# MAJEC: unified gene, isoform, and locus-level transposable element quantification from RNA-seq

**DOI:** 10.64898/2026.04.10.717472

**Authors:** Tian-Yeh Lim, Ari J. Firestone

**Author notes:** To whom correspondence should be addressed. Address: 1130 Veterans Blvd, South San Francisco, CA 94080.

## Abstract

**Background:** The study of transposable elements (TEs) has become increasingly central to fields such as cancer biology, immunology, and aging. Accurately quantifying disease- or laboratory-mediated perturbations in these elements is critical to support this expanding research, yet current RNA-seq pipelines struggle with the pervasive overlap between TEs and protein-coding genes. Existing tools either aggregate to the subfamily level with no locus resolution (TEtranscripts), or provide locus-level quantification without modeling gene overlap (Telescope), with the latter attributing over 40% of TE signal to the 1.1% of loci that overlap gene exons.

**Results:** We present MAJEC (Momentum Accelerated Junction Enhanced Counting), a unified Expectation-Maximization (EM) framework that jointly quantifies genes, transcript isoforms, and individual TE loci from BAM alignments in a single pass. Splice junction evidence informs transcript-level priors, enabling MAJEC to probabilistically distinguish genic from TE-derived reads. This approach was independently validated against Salmon and RSEM on isoform quantification benchmarks. The joint feature space reduces exon-overlap contamination of locus-level TE estimates from 43% of total signal (Telescope) to 5% (MAJEC), while preserving subfamily-level accuracy (differential expression r = 0.987 vs TEtranscripts). Using paired biological vignettes, we demonstrate that MAJEC correctly resolves both the false TE reactivation artifacts endemic to TE-only models, and the false gene upregulation artifacts that occur when heuristic rules misassign genuine intragenic TE transcription.

**Conclusion:** MAJEC simultaneously produces the isoform and locus-level resolution that TEtranscripts lacks, with greater accuracy than Telescope, and runs faster than either.

## 1. Introduction

RNA-seq quantification for genes and their isoforms versus transposable elements (TEs) has bifurcated into two largely independent ecosystems. On the gene side, several distinct method classes coexist: alignment-free tools such as Salmon (Patro *et al*. 2017) and Kallisto (Bray *et al*. 2016) estimate isoform abundance through lightweight sequence-level modeling; alignment-based expectation-maximization frameworks such as RSEM (Li *et al*. 2010; Li and Dewey 2011) resolve multi-mapping reads at the isoform level from aligned reads; and simpler counting tools such as featureCounts (Liao, Smyth, and Shi 2014) and HTSeq (Anders, Pyl, and Huber 2014) tally reads for unambiguous gene-level quantification. For TEs, a separate set of specialized tools has emerged to address the multi-mapping problem inherent in quantifying repetitive sequences: TEtranscripts (Jin *et al*. 2015) distributes multi-mapped reads across TE subfamilies using an expectation-maximization (EM) algorithm, Telescope (Bendall *et al*. 2019) extends this to locus-level resolution with a Bayesian reassignment model and SQuIRE (Yang et al. 2019) adds locus-level TE quantification alongside a separate gene-expression estimate. In practice, obtaining accurate quantification across transcript isoforms, genes, and individual TE loci still requires combining tools that each address only part of the problem, with different alignment requirements, annotation formats, and statistical models; and none resolve gene–TE read overlap within a single model.

Approximately 45% of the human genome derives from transposable elements (International Human Genome Sequencing Consortium, 2001), and interest in TE biology has intensified as roles for TE reactivation have emerged in cancer, aging, innate immunity, and epigenetic therapy (Griffin et al., 2021; De Cecco et al., 2019; Chiappinelli et al., 2015; Roulois et al., 2015; Burns, 2017). A growing number of studies require not just subfamily-level differential expression — “are L1PA7 elements upregulated as a class?” — but locus-level resolution: which *specific* TE insertions are reactivated, where are they in the genome, and what is their relationship to overlapping genes? This granularity is essential for applications such as identifying TE-derived biomarkers of epigenetic drug response, characterizing chimeric TE-gene transcripts, and understanding the regulatory consequences of individual TE insertions.

TEtranscripts, the most widely used TE quantification tool, provides robust subfamily-level differential expression analysis but does not resolve individual TE loci. It also carries a substantial computational cost, which becomes a practical bottleneck for large experiments. Telescope addresses the locus resolution gap but introduces a different limitation: it operates entirely within a TE-only feature space with no access to gene annotations and no modeling of strand information. Since TE insertions are densely embedded within gene bodies — in introns, untranslated regions, and occasionally exons — Telescope has no mechanism to distinguish reads originating from an independently transcribed TE from reads belonging to the host gene’s transcript that happen to overlap a TE annotation. We show that this is not a minor edge case: nearly half of Telescope’s total TE signal originates from exon-overlapping loci, which constitute only 1.1% of all TE features. Thus a 40-fold enrichment for locus-level TE expression is potentially driven by genic signal at exon-TE boundaries.

TEtranscripts mitigates this problem at the subfamily level through a hard-coded rule that assigns uniquely mapped reads overlapping both a gene exon and a TE annotation exclusively to the gene. This conservative heuristic is effective — MAJEC and TEtranscripts agree at r = 0.995 at the subfamily level despite completely different algorithmic approaches. However, the rule does not generalize to locus-level quantification, where the question is not whether a read belongs to “genes” or “TEs” in aggregate but whether a specific read at a specific locus reflects gene or TE transcription. At this resolution, a rigid rule that unconditionally privileges one feature type over the other cannot be correct in both directions: it will suppress genuine TE signal at genic loci just as readily as it suppresses false TE signal.

A joint model that quantifies genes and TEs simultaneously in a shared feature space would resolve this problem by allowing reads to be assigned to whichever feature — gene or TE — is best supported by the data at each locus. Such a model requires accurate isoform-level quantification of the genic component, since reads must be assigned not just to genes but to specific transcripts in order to determine whether they are consistent with the gene’s splice structure or are better explained by independent TE transcription. Splice junction–spanning reads provide particularly strong evidence: a read spanning a known splice junction of a transcript can be confidently assigned to that transcript rather than to an overlapping TE. Junction evidence thus serves a dual purpose — it improves isoform quantification accuracy, and it provides the mechanistic basis for distinguishing genic from TE-derived signal at overlapping loci.

Here we present MAJEC, a unified framework that jointly quantifies genes, transcript isoforms, and individual TE loci from aligned RNA-seq reads (BAM files) in a single analysis. MAJEC operates a locus-level EM algorithm — preserving the individual TE resolution of Telescope — within a joint gene+TE feature space where all annotated transcripts and all annotated TE loci compete for reads simultaneously. Junction-spanning reads inform transcript-level priors through a completeness penalty system that downweights transcripts lacking expected splice junctions, improving isoform accuracy on complex genes and providing high-confidence evidence for genic read assignment at gene-TE overlaps.

We validate MAJEC’s isoform quantification against Salmon on synthetic transcriptomes (Sequins[(Hardwick *et al*. 2016, Dong *et al*. 2023)]) and against both Salmon and RSEM on complex real transcriptomes (LongBench [(You *et al*. 2025)]), demonstrating that junction-informed priors improve accuracy specifically where isoform structure is ambiguous. We show that MAJEC matches TEtranscripts at the subfamily level for differential expression (r = 0.987) while providing locus-level resolution that TEtranscripts cannot, and we demonstrate through paired vignettes that the joint model correctly resolves cases where existing tools fail in opposite directions — Telescope falsely reporting TE reactivation from genic reads, and TEtranscripts falsely inflating gene expression from TE-derived reads. MAJEC completes these analyses faster than the specialized TE tools it replaces, from a BAM alignment that most RNA-seq workflows already produce.

## 2. Methods

### 2.1 Overview

MAJEC quantifies genes, transcript isoforms, and individual TE loci from coordinate-sorted BAM files in a single analysis. The pipeline proceeds in five stages: (1) initial read assignment of uniquely mapping reads via featureCounts, with fractional apportionment when reads overlap multiple annotated features, (2) extraction and scoring of splice junction evidence, (3) featureCounts-based generation of multi-mapper equivalence classes, (4) a two-phase expectation-maximization (EM) algorithm operating on a joint gene+TE feature space, and (5) confidence scoring and output generation.

MAJEC requires as input a coordinate-sorted BAM file from any splice-aware aligner (e.g., STAR (Dobin *et al*. 2012) or HISAT2), a gene annotation GTF (Ensembl or GENCODE), and a TE annotation file (RepeatMasker (Smit, Hubley, and Green 2013)). For TE quantification, the upstream aligner must retain multi-mapping reads (e.g., STAR with or higher); this is the same requirement as TEtranscripts and Telescope. MAJEC runs on standard Linux workstations with 32–64 GB RAM and does not require HPC infrastructure.

### 2.2 Feature annotation and joint model design

MAJEC constructs a unified feature space containing all annotated transcript isoforms from the gene GTF and all annotated TE loci from RepeatMasker. In this joint model, a read overlapping both a gene exon and a TE annotation competes between the two feature types — the EM assigns it to whichever feature is best supported by the available evidence. This is the key architectural difference from existing TE tools.

The EM operates at individual transcript and TE locus resolution — preserving the locus-level granularity. Subfamily-level and gene-level counts are derived by post-hoc aggregation of locus-level estimates.

Subset and superset relationships between transcripts are identified during annotation preprocessing. A transcript whose set of annotated splice junctions is entirely contained within another transcript’s junction set is flagged as a potential subset isoform, subject to the subset penalty system described below.

To accurately apply these penalties, MAJEC employs an annotation preprocessing step. The pipeline constructs a deterministic index of all expected splice junctions, categorizing each as either unique to a specific isoform or as a site of direct splice competition (e.g., shared donor but alternative acceptor sites). Furthermore, rather than uniformly penalizing subset transcripts, MAJEC utilizes interval-tree logic to identify “unambiguous exonic territories”—genomic coordinates strictly unique to the subset isoform. By categorizing the structural basis of these unique regions (such as retained introns or alternative terminal extremities), the pipeline extracts isolated regions to serve as “rescue features.” During quantification, coverage across these precise unique intervals can empirically validate a subset’s expression, rescuing it from being erroneously zeroed out by its superset.

### 2.3 Initial read assignment

Equivalence classes are built from two sequential featureCounts runs. Initial counts for reads with no mapping ambiguity are collected first. After filtering the BAM for multi-mapped reads, a second set of equivalence classes is generated with fractional counting enabled for reads mapping to multiple features. MAJEC modifies the default featureCounts fractional counting behavior by pooling all features across all alignment positions for each multi-mapped read, distributing the count equally across the union of overlapping features rather than independently at each alignment position. This ensures that the initial count for a multi-mapper reflects the full set of candidate features before EM resolution. Unlike the default multi-overlap mode in featureCounts, MAJEC’s union-based fractional assignment strictly conserves total read mass: each read contributes exactly one count distributed across all candidate features. For paired-end data, counting is performed at the fragment level, and strandedness is handled according to the library preparation protocol. To ensure memory efficiency, the featureCounts processes are streamed via FIFO pipes, allowing for the concurrent processing of multiple BAM files.

### 2.4 Junction evidence and prior adjustments

Before the EM begins, initial counts from featureCounts are modulated by a series of evidence-based priors computed from splice junction observations. Junction-spanning reads are extracted from the BAM file and mapped to annotated splice sites in the GTF, providing per-sample evidence for which transcripts are actively spliced.

Four prior adjustments are applied sequentially:

#### Junction evidence boost

Transcripts receive additional counts proportional to their splice junction support, weighted by junction uniqueness. For a junction *j* shared by *n* transcripts, the evidence contributed to each transcript is scaled by (1/*n*)^*d*, where *d* is a decay exponent (default 1.0). This ensures that junctions unique to a single transcript provide strong evidence, while shared junctions contribute proportionally less. The magnitude of the boost is controlled by a global junction weight parameter (default 2.0).

#### Transcript Support Level (TSL) penalty

Transcripts are optionally down-weighted based on their GENCODE/Ensembl Transcript Support Level annotation, ranging from (default = 1.0 (TSL1, highest confidence) to 0.3 (TSL5, lowest confidence), with 0.8 for transcripts with no TSL annotation). Use and weighting of TSL-based penalties are user configurable.

#### Junction completeness penalty

For each multi-exon transcript, MAJEC compares observed junction-spanning read counts to the expected coverage based on the transcript’s median junction depth. Junctions with substantially lower coverage than expected receive a Z-score-based penalty, incorporating an overdispersion parameter (default 0.05) that models biological and technical variance. Terminal junctions (5’ or 3’ ends) receive relaxed penalties when appropriate for the library type (e.g., 3’ dropout in poly-dT libraries). The completeness penalty ranges from 1.0 (all expected junctions observed at expected coverage) to a configurable minimum (default 0.0001). This penalty is the primary driver of MAJEC’s isoform accuracy advantage, as demonstrated by feature ablation (see Results).

#### Subset isoform penalty

Transcripts identified as potential subsets of longer isoforms during annotation preprocessing are evaluated in two stages. First, an annotation-based penalty compares the subset’s shared junction coverage to the expected contribution from its superset transcripts; if the observed coverage does not exceed the expected superset contribution (controlled by a threshold parameter, default 1.25), an initial penalty is applied. Second, when unique exonic territory data is available, direct read coverage in the subset’s unique regions is compared to coverage in shared regions, providing data-driven validation or override of the annotation-based penalty. The subset penalty floor (default 0.001) controls the extent of subset transcripts suppression when the evidence is ambiguous.

The adjusted counts after all four prior stages serve as the initial estimates for the EM algorithm.

### 2.5 Two-phase EM algorithm

MAJEC employs a hierarchical two-phase EM that processes unique and multi-mapping reads separately, recognizing that they represent fundamentally different levels of uncertainty.

In Phase 1, uniquely mapped reads are assigned to transcripts and TE loci based on the prior-adjusted initial estimates. Because these reads have unambiguous genomic locations, the only ambiguity is among overlapping features at the same locus (e.g., overlapping isoforms, or a gene exon overlapping a TE annotation). In Phase 2, multi-mapping reads are iteratively allocated using a combination of the fixed unique-mapper estimates from Phase 1 and the updated multi-mapper probabilities from the preceding EM iteration.

The E-step allocates reads within each equivalence class (a group of reads sharing the same set of possible feature assignments) proportionally to current expression estimates:

For transcript *t* in equivalence class *C*:

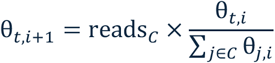

The combined estimate after both phases is θ_total,*t*,*i*+1_ = θ_unique,*t*,*i*+1_ + θ_multi,*t*,*i*+1_.

#### Momentum acceleration

MAJEC accelerates convergence using expression-level grouped momentum. After an initial burn-in period, velocity vectors (θ[*t*, *i*] − θ[*t*, *i*−1]) are computed per transcript and scaled by expression-group-specific momentum factors (default: 1.5 for low-expression, 1.0 for medium, 0.7 for high-expression transcripts). Stability safeguards include oscillation detection, maximum step-size limits, and automatic fallback to standard EM upon detection of instability.

#### Convergence

The EM iterates until relative change, absolute change, and total count stability all fall below adaptive thresholds, or until the iteration cap is reached (default 100). In practice, convergence occurred within 15–17 iterations across all tested samples; the iteration cap is a safety net rather than a tuning parameter.

#### Effective length correction

Transcript effective lengths are computed from the empirical fragment length distribution (estimated from the BAM file or provided by the user) using a Normal distribution model of mappable positions along each transcript.

### 2.6 Confidence metrics

MAJEC reports several per-transcript and per-group confidence metrics alongside the count estimates.

#### Distinguishability score

A composite metric combining the fraction of reads with unambiguous assignment (unique mappers plus unique junction evidence from multi-mappers) with a measure of how well the transcript was separated from competitors within the ambiguous read fraction (ambiguous_distinguishability):

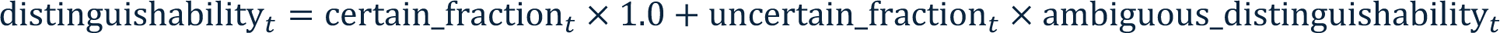

Scores range from 1.0 (perfectly distinguishable) to 0.0 (highly ambiguous).

#### Assignment entropy

Shannon entropy of the read source distribution for each transcript, converted to a confidence score: confidence[*t*] = 1 / (1 + entropy[*t*]).

#### Discord score

A measure of EM revision magnitude, computed as an expression-normalized log reduction from pre-EM to post-EM counts. For each transcript with pre-EM count *p* > post-EM count *q*:

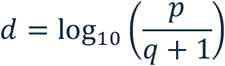

This raw discord is normalized by an expression-dependent scale factor 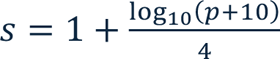, ensuring that proportionally equivalent revisions receive similar scores regardless of expression level. A saturating transform bounds the final score on [0, 4):

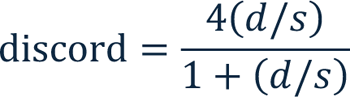

Transcripts where the EM did not reduce counts receive a score of zero. High discord combined with unique junction support identifies a small population (∼2% of high-discord transcripts) representing potential EM over-suppression candidates.

#### Group-level metrics

For genes and TE subfamilies, expression-weighted averages of transcript-level scores are computed, along with an inter-group competition metric that distinguishes ambiguity from complex internal splicing (low inter-group competition) versus ambiguity from external paralogs or gene-TE overlap (high inter-group competition).

### 2.7 Output

MAJEC produces transcript-level and TE locus-level count estimates, which are aggregated to gene-level and TE subfamily-level counts by summation. MAJEC outputs transcript-level and TE locus-level count estimates as tab-delimited files. Optional companion utilities ingest these results into a SQLite database for downstream querying, extract DESeq2-compatible count matrices at the gene, isoform, and TE locus levels, and generate interactive HTML reports (Plotly-based) for isoform-level visualization of junction evidence, expression proportions, and confidence metrics.

### 2.8 Benchmarking datasets

#### 2.8.1 Sequins v2.4 (synthetic isoforms)

The Sequins v2.4 spike-in dataset comprises 160 synthetic RNA isoforms across 76 genes with known ground truth concentrations in two mixes (Mix A and Mix B), spiked into human RNA and sequenced as paired-end short reads. Reads pairs (38.9–61.6 M) were aligned with STAR 2.7.11b to a hybrid hg38 + Sequin chromosome genome using--outFilterMultimapNmax 100 and gene annotations from refGene (appropriate for the synthetic transcript models). MAJEC, Salmon 1.10.3 (with --gcBias –seqBias) featureCounts 2.1.1 (raw fractional counts, pre-EM), and long-read quantification were compared. Accuracy was assessed using median absolute log2 fold error (with pseudocount of 1) and Pearson correlation in log-transformed space.

#### 2.8.2 LongBench / ENCODE (real transcriptomes)

Eight ENCODE lung cancer cell lines (SHP-77, NCI-H146, NCI-H526, NCI-H2228, NCI-H69, HCC827, NCI-H211, NCI-H1975) with matched short-read (55–71 M read-pairs) and long-read data were used for isoform quantification benchmarking. Short reads were aligned with STAR 2.7.11b using --outFilterMultimapNmax 100 against the GENCODE v44 annotation (Frankish *et al*. 2020). Long-read data from three platforms — ONT direct RNA (dRNA), PacBio Iso-Seq, and ONT cDNA — were aligned with pbmm2 1.9.0 (--preset ISOSEQ) and quantified with IsoQuant 3.9.0 to serve as approximate ground truth, along with their consensus average (mean across platforms). MAJEC, Salmon 1.10.3 (with --gcBias -- seqBias), and RSEM 1.2.28 were compared.

Transcripts were normalized to counts-per-10-million and per-transcript quantification error was computed as |log1p(observed) - log1p(truth) |. The primary comparison metric was the superiority rate: the fraction of transcripts where MAJEC had lower error than the comparator, with ties split equally. Statistical significance was assessed by Wilcoxon signed-rank test on paired error differences. Analyses were stratified by MAJEC’s junction penalty classification (none, incomplete, subset, or both penalties below 0.2), by ground truth expression level, and by transcript detection (sensitivity, precision, F1).

Intrinsic transcript properties analysis was conducted on per-transcript accuracy outcomes using the full LongBench benchmark modeled against intrinsic transcript features. The analysis was restricted to protein-coding and lncRNA transcripts using the GENCODE v44 biotypes. For each transcript in each cell line, a “win” was using the superiority metric and error calculated as above against the consensus long-read ground truth (mean of PacBio, ONT, and dRNA) on CPM normalized to 10 M. Intrinsic features (isoforms per gene, exon count, spliced transcript length, and GC content) were computed from the GENCODE v44 annotation and the matching human genome, with log10 expression (CPM) as a covariate. Feature effects were estimated by multivariable logistic regression on decided cases (exact ties excluded) with all predictors standardized (variance inflation factors < 1.8), and by ordinary least squares on the continuous error difference; standard errors were made robust by clustering on gene (30,869 clusters) to account for transcripts observed across the eight cell lines.

For sequencing depth analysis, 3 lines (SHP-77, NCI-H2228, NCI-H146) were downsampled to 7.5, 15, 30, and 55 M read pairs with seqtk 1.5-r133 (Li, 2025; fixed seed, applied identically to both mates). Each subset was aligned with STAR and quantified with both MAJEC and Salmon as above from the identical read subset. Isoforms were classified by their long-read intra-gene structure as dominant, minor, or rare (rare = a minor isoform contributing <5% of its gene’s long-read signal), and median log-fold error and detection sensitivity using a threshold of 1 in the per-10-million–normalized count were computed on the long-read– confirmed isoforms within each stratum.

An independent replication was performed on a T-cell dataset GSE229972 (Woolley et al., 2025) with matched short-read (58.3–69.4 M read-pairs) and PacBio Iso-Seq data from three timepoints. Short reads were aligned with STAR 2.7.11b against GENCODE v29 annotations; PacBio reads were aligned with pbmm2 1.9.0 (Biosciences, n.d.), a PacBio-native frontend for the minimap2 (Li H 2018) and quantified with IsoQuant 3.9.0. (Prjibelski *et al*. 2023).

#### 2.8.3 TE benchmarking (decitabine treatment)

Stranded paired-end RNA-seq (28.0–35.5 M read-pairs per sample) was generated from FaDu (ATCC, HTB-43) head and neck squamous carcinoma cells treated with DMSO or 1 µM decitabine for 7-days in triplicate. Reads were aligned with STAR 2.6.1a with --outFilterMultimapNmax 100 --winAnchorMultimapNmax 200, retaining the alignment used for the original TEtranscripts analysis. Gene annotations were from refGene (hg38refGene); TE annotations were from RepeatMasker (hg38_rmsk_TE, locus-level). MAJEC was run in two modes: joint (gene+TE feature space) and TE-only (identical EM without gene features). Results were compared to SQuIRE 0.9.9.92, TEtranscripts 2.2.3, Telescope 1.0.3 (--reassign_mode conf), and an additional development branch of Telescope with a stranded mode (1.0.3+48.g4cf1859) run with the same command plus --stranded_mode RF. MAJEC, TEtranscripts, and both Telescope builds used the shared STAR 2.6.1a alignment. SQuIRE was instead run through its native Fetch/Clean/Map/Count pipeline, which performs its own STAR 2.5.3a alignment (--read_length 150, default multi-mapping); because SQuIRE Map aligns with --alignEndsType EndToEnd, reads for SQuIRE were first adapter- and quality-trimmed with Trim Galore 0.6.10 (Krueger, 2023; Cutadapt 1.18, Martin 2011); paired-end, Phred 20, minimum length 20 bp, Illumina adapter auto-detected). Subfamily-level comparisons used Pearson correlation and DESeq2 differential expression concordance (log2 fold change correlation, significance concordance). Locus-level comparisons used per-locus count ratios (log2(MAJEC/Telescope)) stratified by overlap classification.

##### Overlap classification

TE loci were classified as intergenic, intronic, or exon-overlapping based on their intersection with annotated gene features. To process the ∼4.7 million annotated TE loci efficiently, classification was performed using a custom, chromosome-parallelized Python script utilizing a binary search algorithm over sorted genomic intervals. A TE locus was first evaluated for overlap with any annotated gene body; loci lacking overlap were classified as intergenic. Genic TEs were subsequently evaluated against exon intervals. To match Telescope’s unstranded quantification model, this exon intersection was performed independent of strand direction. Loci overlapping an exon on either strand were classified as exon-overlapping, while the remaining genic loci were classified as intronic. Of 4.69 million annotated TE loci, 54% were intergenic, 45% intronic, and 1.1% exon-overlapping.

##### Differential expression comparison

DESeq2 1.38.3 (Love, Huber, and Anders 2014) was run independently on MAJEC and Telescope count matrices. For MAJEC, the full count matrix (gene transcripts + TE loci) was provided to DESeq2, with genes contributing to size factor estimation; for Telescope, TE loci only were provided, consistent with each tool’s standard usage. TE loci with ≥5 total counts in ≥2 sampels for a given tool were retained for comparison. Significance was assessed at padj < 0.05.

##### Vignette analysis

Individual loci exhibiting discordant calls between tools were examined using stranded coverage plots (fractional depth, 1/NH weighting) generated from the BAM files. For the TEtranscripts comparison, DESeq2 was run on TEtranscripts output and compared to MAJEC gene-level and TE locus-level results at loci where MAJEC identified TE-dominated signal (TE log2FC > gene log2FC).

##### Gene-only (TE-free) simulation

Reads were simulated with polyester 1.38.0 (Frazee et al., 2015) exclusively from GENCODE v44 protein-coding and lincRNA transcripts, with no transposable-element sequence present (40,001,333 fragments per sample, two replicates, 150 bp, error rate 0.005). Any read a quantifier assigns to a TE locus is therefore a false positive by construction. Simulated reads were aligned and quantified with the same STAR parameters and MAJEC, released Telescope (v1.0.3, unstranded), and development-branch Telescope (1.0.3+48.g4cf1859, stranded) settings as the decitabine benchmark. False-positive rate was computed as the fraction of simulated signal assigned to TE loci, and the residual false positives were broken down by TE family.

#### 2.8.4 Parameter sensitivity and feature ablation

Forty-three parameter conditions were evaluated on the LongBench dataset using MAJEC’s cached mode, which reuses featureCounts assignments and junction metrics across runs (∼14 minutes per condition, ∼11 hours total CPU). Feature ablation removed each prior component individually (completeness penalties, subset penalties, junction boosts, paired rescue, terminal recovery, territory adjustments). Parameter sweeps varied individual parameters across their range while holding others at defaults. Performance was assessed as median log1p fold error against consensus long-read ground truth, with Wilcoxon signed-rank tests comparing each condition to Salmon.

### 2.9 Software and data availability

MAJEC v0.1.0 is implemented in Python and is available at https://github.com/calico/majec under the Apache 2.0 license. All analyses used the following software: STAR 2.7.11b (isoform benchmarks), STAR 2.5.3a (SQuIRE), and 2.6.1a (TE benchmark), Salmon 1.10.3, RSEM 1.2.28, Telescope 1.0.3, TEtranscripts 2.2.3, SQuIRE 0.9.9.92, DESeq2 1.38.3, featureCounts 2.1.1 (Subread), seqtk 1.5-r133, polyester 1.38.0, Trim Galore 0.6.10, Cutadapt 1.18, IsoQuant 3.9.0, and pbmm2 1.9.0. Gene annotations were from GENCODE v44 (LongBench), GENCODE v29 (T-cell), and refGene (Sequins, TE benchmark); TE annotations were from RepeatMasker (hg38). Notebooks and data for figure generation are located at https://doi.org/10.5281/zenodo.19224158. RNA-seq data generated for this study are deposited at SRA PRJNA1454087. LongBench data were obtained from GSE303762. Raw short read Sequins spike-in data were obtained from GSE164598 and sequins GTF, ground truth, and processed longread data are from https://github.com/XueyiDong/LongReadRNA.

## 3. Results

### Junction-informed isoform quantification is accurate and mechanistically grounded

The joint gene+TE model depends on accurate assignment of genic reads to their correct transcripts — errors in isoform quantification propagate directly into TE estimates. We therefore first validated MAJEC’s junction-informed EM (Fig. 1A) against established isoform quantification tools using two complementary benchmarks.

**Figure 1.**
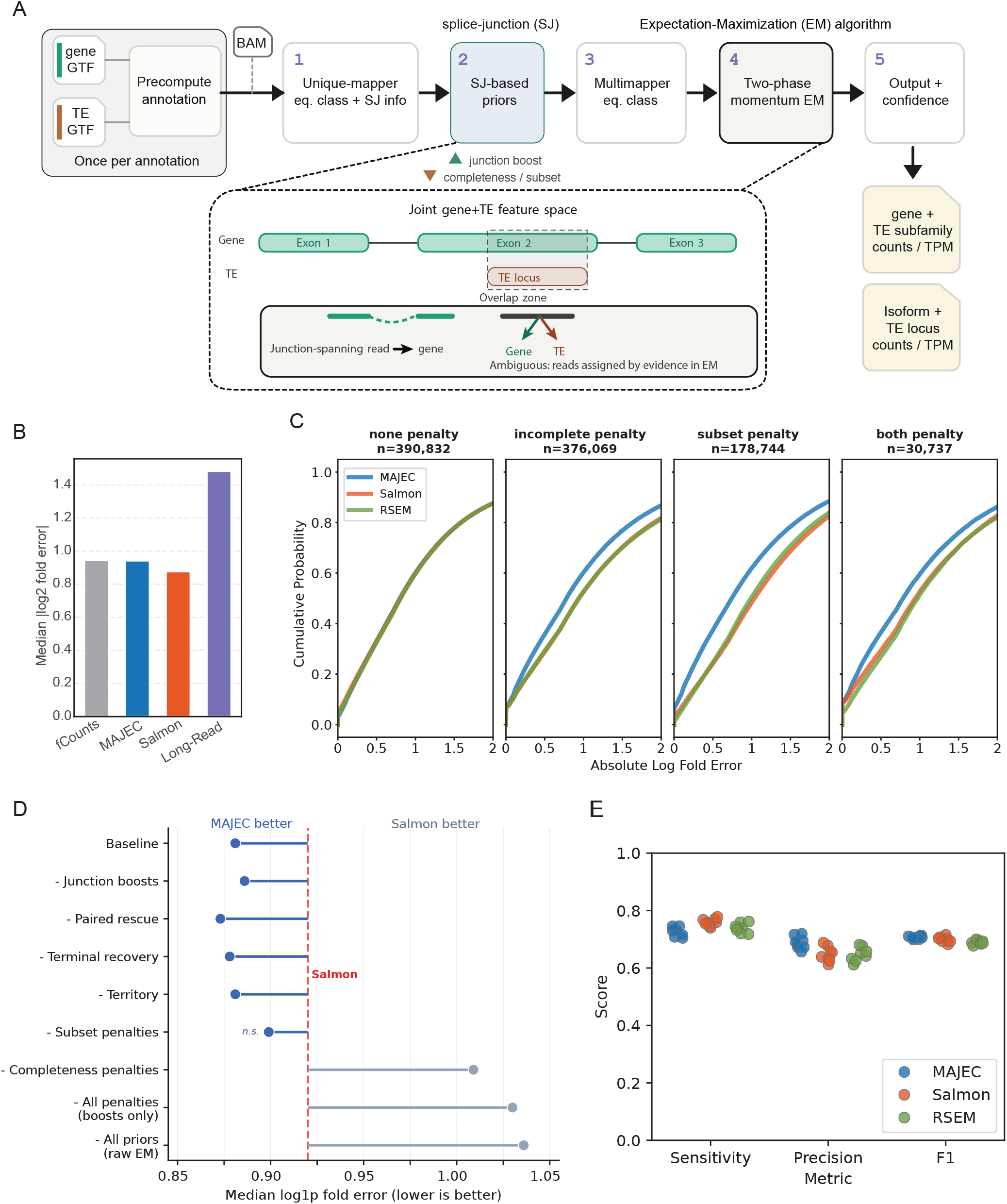
MAJEC isoform quantification benchmarking. **(A)** Schematic depicting information flow and design of the MAJEC quantification pipeline. From the gene and TE annotations, a shared feature index is precomputed once per annotation (grey). For each sample MAJEC then (1) builds equivalence classes (“eq. class”) for uniquely mapped reads and extracts splice-junction (SJ) evidence from read alignments; (2) converts that SJ evidence into transcript-level priors, boosting isoforms whose annotated junctions are observed and penalizing those with missing or under-covered junctions; (3) builds equivalence classes for multi-mapping reads; (4) resolves all reads with a two-phase, momentum-accelerated expectation–maximization (EM) algorithm that fixes uniquely mapped reads first and then allocates multi-mappers; and (5) reports counts (and TPM) at gene + TE-subfamily and isoform + TE-locus levels with per-feature confidence. *Inset: Joint gene+TE feature space.* A read falling in an overlap zone (here a TE locus within exon 2) MAJEC lets the two features compete for each such read in the EM, weighted by the SJ-derived priors and other unambiguous evidence. **(B)** Median log2 fold error against known ground truth concentrations for the Sequins v2.4 spike-in dataset (160 isoforms across 76 genes). **(C)** CDF of log fold error of MAJEC, Salmon, RSEM against consensus long-read ground truth (mean of dRNA, PacBio Iso-Seq, and ONT cDNA). Pooled transcript measurements from 8 different cell lines are analyzed and stratified by the junction-based penalty assigned by MAJEC (see Methods). MAJEC performance is best on penalized transcripts. **(D)** Feature ablation analysis. Each row shows the effect of disabling a single MAJEC feature on median log1p fold error across 8 cell lines relative to the consensus long read data. The dashed red line indicates Salmon’s median error. Subset and completeness penalties contribute the largest individual improvements. Removing subset penalties alone results in a statistical tie with Salmon (Wilcoxon signed-rank test, p > 0.05). “All penalties” removes completeness and subset penalties while retaining junction boosts; “All priors” disables all junction-informed adjustments, leaving only the unweighted EM. **(E)** Sensitivity, precision, and F1 for each short read quantification against the consensus long-read values. Data from the eight individual cell lines are represented as dots.

On the Sequins v2.4 synthetic spike-in dataset (160 isoforms across 76 genes with known ground truth concentrations), MAJEC, Salmon, and raw featureCounts fractional counting all achieved similar accuracy, with median absolute log2 fold errors of 0.88–0.95 (Fig. 1B, S1). No method was meaningfully better or worse on this simple transcriptome (max 4 isoforms per gene) where all annotated transcripts are explicitly present, confirming that MAJEC’s junction priors do not introduce systematic bias when they have little work to do.

On complex real transcriptomes — eight ENCODE lung cancer cell lines with matched short-read and long-read data (LongBench) — a different picture emerged. Pooling across cell lines and long-read ground truths (∼1 million transcript-level comparisons), MAJEC produced lower quantification error than Salmon on 54.3% of transcripts (median Δ = −0.041, Wilcoxon p < 2.2 × 10⁻³⁰⁸) and lower error than RSEM on 53.7% of transcripts (median Δ = −0.039, p < 2.2 × 10⁻³⁰⁸). The advantage was consistent across all eight cell lines and all four long-read ground truths (dRNA, PacBio Iso-Seq, ONT cDNA, and their consensus average), with the dRNA platform showing the largest MAJEC advantage (Δ = −0.079 vs. Salmon). In addition to error reduction, MAJEC also demonstrated modest improvements in Pearson correlation across all cell lines and ground truths (Fig. S2).

To isolate the mechanism driving this improvement, we stratified transcripts based on the two junction-based priors MAJEC computes prior to the EM: a completeness penalty (reflecting whether expected splice junctions were missing) and a subset penalty (reflecting whether the transcript’s junction evidence is better explained by an overlapping isoform). Transcripts can receive either penalty, both, or neither (Fig. S3). Transcripts with strong, isoform-specific junction support also receive a positive boost to their initial estimates.

Stratifying the data by these junction penalty classifications revealed a clear dependence between MAJEC’s performance advantage and structural complexity (Fig. 1C). Transcripts with minimal junction-based penalties (∼40% of the transcriptome) showed no MAJEC advantage — all three methods performed equivalently. Transcripts penalized for incomplete junction coverage (∼37%) showed a clear MAJEC advantage (57.0% win rate, Δ = −0.102), and transcripts penalized as subsets of better-supported isoforms (∼17%) showed the largest improvement (62.3% win rate, Δ = −0.167). This establishes that MAJEC’s advantage originates directly from its junction-informed initial estimates: transcripts entering the EM with strongly penalized starting points achieve significantly improved accuracy, while those with unpenalized starting points maintain baseline parity with competing methods. These results generalized well across all individual cell lines and ground truths (Fig. S2).

Feature ablation on the LongBench dataset confirmed this interpretation from the opposite direction (Fig. 1D). Removing completeness penalties — the largest single contributor — caused MAJEC to lose to Salmon by approximately 10% in median error. Removing subset penalties reduced the advantage to a statistical tie (p = 1). Removing both penalty types further degrade model performance, demonstrating that the core innovation is penalizing implausible transcript assignments rather than boosting plausible ones. The remaining features (paired rescue, terminal recovery, territory adjustments) had marginal global impact on this dataset (Fig. S4).

To further explore what drives the modest difference between MAJEC and Salmon, we tested whether intrinsic transcript properties predict which tool is more accurate. Restricting to protein-coding and lncRNA transcripts (96% of the annotated set; the biotypes reliably measured by long-read sequencing), we modeled per-transcript win/loss as a function of isoform count, exon count, spliced length, GC content, and expression in a standardized multivariable logistic regression (n = 969,837 transcript–sample observations, S5A). MAJEC’s advantage concentrated on structurally complex, low-abundance transcripts: the win rate rose with isoform count (49.7% for single-isoform genes to 60.0% for genes with ≥6 isoforms; OR 1.10 per standard deviation [SD]), peaked at intermediate exon counts (58.0% at 4–6 exons; OR 1.06 per SD), and was highest at low expression (62.0% in the lowest expression quintile, falling to 42.1% in the highest; OR 0.89 per SD; Fig. S5). The performance of MAJEC is thus relatively strongest in the regimes in which read-to-isoform assignment is most ambiguous and junction-informed priors are most informative. Conversely, where there is less structural ambiguity to resolve, the two tools were near parity: single-isoform genes (49.7%) and single-exon transcripts (48.5%; S5B). Salmon’s clearest advantage was on longer, more highly expressed transcripts (spliced length OR 0.74 per SD), where read evidence is abundant and assignment is largely unambiguous. GC content had no independent effect on the outcome (OR 1.00).

At the transcript detection level, MAJEC traded modest sensitivity for improved precision (Fig. 1E). Compared to Salmon, MAJEC detected fewer true transcripts (sensitivity 0.727 vs. 0.759) but called 7,000– 9,000 fewer false positives per cell line (precision 0.688 vs. 0.648), resulting in a higher composite F1 score (0.706 vs. 0.699 for Salmon; 0.690 for RSEM). This precision advantage is directly relevant to the joint gene+TE model: suppressing unsupported gene transcripts prevents false genic features from competing with TE loci for shared reads during the EM.

These findings replicated on an independent T-cell dataset, where MAJEC showed the same penalty-type-dependent advantage over Salmon and RSEM, with a smaller but consistent effect size (50.9% win rate vs RSEM; full results in Fig. S6).

To examine how sequencing depth affects accuracy for MAJEC, we downsampled the short reads of three LongBench cell lines to 7.5, 15, 30, and 55 M read pairs per line and re-quantified each subset with MAJEC and Salmon. Isoforms were stratified by their long-read intra-gene structure into dominant, minor, and rare classes (rare = minor isoforms contributing <5% of their gene’s long-read signal), and median log-fold error and detection sensitivity were evaluated on the long-read–confirmed isoforms in each class (Fig. S7). Both tools improved in accuracy with depth. MAJEC’s advantage was concentrated in the rare class: its median error was lower than Salmon at every depth (0.953 → 0.797 from 7.5 to 55 M, versus Salmon 1.071 → 0.924), and MAJEC had the lower error in all three cell lines at every depth. The two tools also differed sharply in the depth dependence of rare-isoform detection: MAJEC’s sensitivity was essentially flat across the range (∼0.70), whereas Salmon’s rose steeply from 0.555 at 7.5 M, drawing level near 30 M and marginally surpassing MAJEC (0.73 vs 0.70) by 55 M. For the two higher-abundance classes the pattern was different: MAJEC held a modest median error edge at low and intermediate depth, but its accuracy was surpassed by Salmon at 55 M read pairs. MAJEC thus shows no tool-specific, disproportionate degradation on low-coverage data, and is if anything comparatively more sensitive and accurate than Salmon when reads are sparse.

Long-read quantification serves as an approximate rather than absolute ground truth, and itself showed the highest error on the Sequins benchmark (median 1.48 vs. 0.88–0.95 for short-read methods). However, several features of the LongBench results argue against the advantage being an artifact of ground truth bias: the improvement was consistent across three independent long-read platforms with distinct error profiles (dRNA, PacBio Iso-Seq, ONT cDNA), was mechanistically confined to transcripts receiving strong junction penalties, and was eliminated by feature ablation. The Sequins and LongBench benchmarks are thus complementary — Sequins validates accuracy against absolute ground truth on simple genes, while LongBench tests relative performance on the complex isoform structures where junction priors are expected to help, using the best available approximate ground truth.

MAJEC’s junction-informed EM has identifiable limitations. On the Sequins dataset, asymmetric isoform pairs with highly imbalanced unique evidence can trigger over-concentration, where the EM iteratively assigns all shared reads to the dominant isoform e.g., R2_7_1(Fig. S1). MAJEC’s advantage over Salmon is strongest at low expression (<3 CPM, 61.4% win rate), where junction penalties effectively suppress marginal transcripts. At moderate expression levels (10–316 CPM), Salmon’s sequence-level modeling — including GC and fragment-length bias correction that MAJEC does not implement — provides an edge (win rates 39–46%). The two methods converge at high expression (>316 CPM), where both have abundant evidence and the modeling differences matter less (Fig. S2C).

### MAJEC produces TEtranscripts-equivalent subfamily quantification

Having established the reliability of MAJEC’s isoform quantification, we next evaluated its TE subfamily estimates against the most widely used TE quantification tool, TEtranscripts, and the locus-resolution tool Telescope. To evaluate TE quantification in a biologically relevant context, we generated stranded 150 bp paired-end RNA-seq from FaDu head and neck squamous cell carcinoma (HNSCC) cells treated with DMSO or 1 µM decitabine for 7-days in triplicate. Decitabine, a DNA methyltransferase inhibitor, induces widespread TE demethylation and reactivation alongside gene-level expression changes, providing a dataset where both gene and TE differential expression are expected — an ideal test case for evaluating gene-TE signal separation undefined.

At the subfamily level, MAJEC and TEtranscripts produced near-identical count estimates (Pearson r = 0.993–0.995 across samples; Fig. 2A, left panel), with strong agreement across all major TE classes (LINE, SINE, LTR, DNA; r = 0.990–0.999; Fig. S8A). This agreement is notable because the two tools use fundamentally different EM architectures: TEtranscripts distributes multi-mapped reads across TE subfamilies directly, while MAJEC resolves reads to individual TE loci and then aggregates to the subfamily level. The convergence of these two approaches on the same subfamily totals provides independent validation of both.

**Figure 2.**
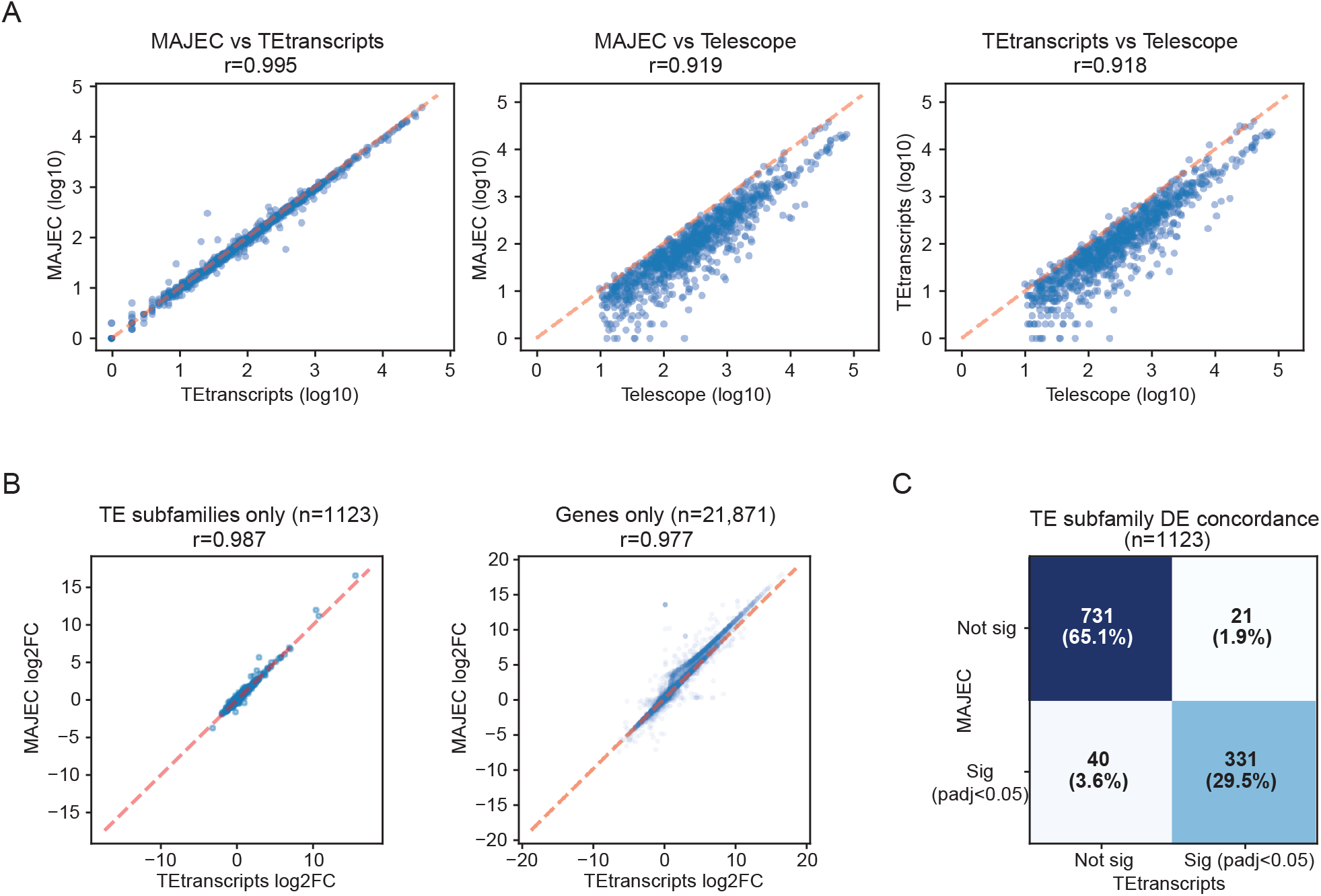
Mechanistically distinct gene-aware TE quantification platforms agree and contrast with TE-only feature space quantification. **(A)** Pair-wise scatter plots showing the agreement of subfamily TE counts from MAJEC, TEtranscripts and Telescope in Log 10 count space. One replicate of the decitabine treated cells are shown but are highly representative of general results. MAJEC and TEtranscripts demonstrate very strong agreement while both similarly diverge from the gene-unaware Telescope data. **(B)** Scatter plots showing Log2 fold change concordance between MAJEC and TEtranscripts for TE subfamilies and genes. **(C)** Concordance matrix showing strong agreement in significance calls by MAJEC and TEtranscripts on subfamilies.

Both MAJEC and TEtranscripts diverged from Telescope by a similar magnitude (MAJEC–Telescope r = 0.919; TEtranscripts–Telescope r = 0.918; Fig. 2A, center and right panels). This symmetric pattern is informative. MAJEC and Telescope use more similar EM architecture — locus-level resolution followed by aggregation — yet MAJEC agrees far more closely with TEtranscripts than with Telescope. The variable that distinguishes the grouping is not the EM granularity but how each tool handles genic contamination. MAJEC addresses it dynamically through its joint gene+TE EM model, where genes and TE loci compete for reads probabilistically. TEtranscripts addresses it through a different mechanism: a hard-coded prioritization rule that assigns uniquely mapped reads overlapping both a gene exon and a TE annotation exclusively to the gene. Telescope, by contrast, operates entirely blind to the gene feature space and retains genic reads with no mechanism to redistribute them. This convergence — a probabilistic joint model and a rule-based heuristic arriving at the same subfamily estimates through completely different strategies — is among the strongest evidence that the joint model is producing correct TE estimates rather than merely different ones.

MAJEC’s EM also improved subfamily estimates relative to the pre-EM fractional counts. The mean absolute log2 fold change between MAJEC and TEtranscripts decreased from 0.40 (pre-EM) to 0.19 (post-EM), indicating that the EM redistributes multi-mapped reads in a manner consistent with TEtranscripts’ subfamily-level model. Notably, the EM’s effect on subfamily estimates was nearly unidirectional, reducing counts as genic reads were reassigned from TE subfamilies to overlapping gene features (Fig. S8B).

At the differential expression level, MAJEC and TEtranscripts showed high concordance. DESeq2 log2 fold change estimates correlated at r = 0.977 overall and r = 0.987 for TE subfamilies specifically (Fig. 2B). Of TE subfamilies called as significantly differentially expressed by either tool, 94.6% were concordant in direction and significance, while the 61 discordant subfamilies (5.4%) still demonstrated strong log2fold correlation (r = 0.88) (Fig. 2C & Fig. S8C). MAJEC can therefore serve as a drop-in replacement for TEtranscripts in existing DE workflows, with the additional benefit of locus-level resolution that TEtranscripts does not provide.

### Joint gene–TE modeling eliminates false TE signal from genic loci

The central motivation for MAJEC’s joint model is the pervasive overlap between TE annotations and protein-coding genes. Approximately 45% of the human genome derives from transposable elements, and many TE loci reside within gene bodies — in introns, UTRs, and occasionally exons. When a gene is transcribed, reads originating from the gene’s mRNA will overlap these embedded TE annotations. A TE-only quantification model has no mechanism to distinguish such genic reads from reads produced by independent TE transcription, leading to systematic false TE signal at gene-overlapping loci.

To quantify this problem, we classified TE loci by their relationship to gene annotations — ignoring strand, since Telescope does not model strandedness — as intergenic (54.2% of loci), intronic (44.7%), or exon-overlapping (1.1%), and measured the fraction of total TE signal originating from each class.

The results revealed exonic overlap to be a dominant source of contamination. In Telescope, exon-overlapping TE loci accounted for 43% of total TE signal (876K of 2.03M fragments; Fig. 3A). This was not issue of strandedness as the unreleased, development, version of Telescope which has a stranded mode left the pattern essentially unchanged (41% exon-overlapping). Moreover, MAJEC run in a TE-only mode (identical EM but without gene features) reproduced it closely (42%) and the same contamination arose in SQuIRE (42%), a tool that *does* incorporate a gene annotation, but quantifies TE loci in a separate feature space. Only MAJEC’s joint model, in which genes and TE loci compete for each read within a single EM, reduced exon-overlapping signal to 5% (Fig. 3A).

**Figure 3.**
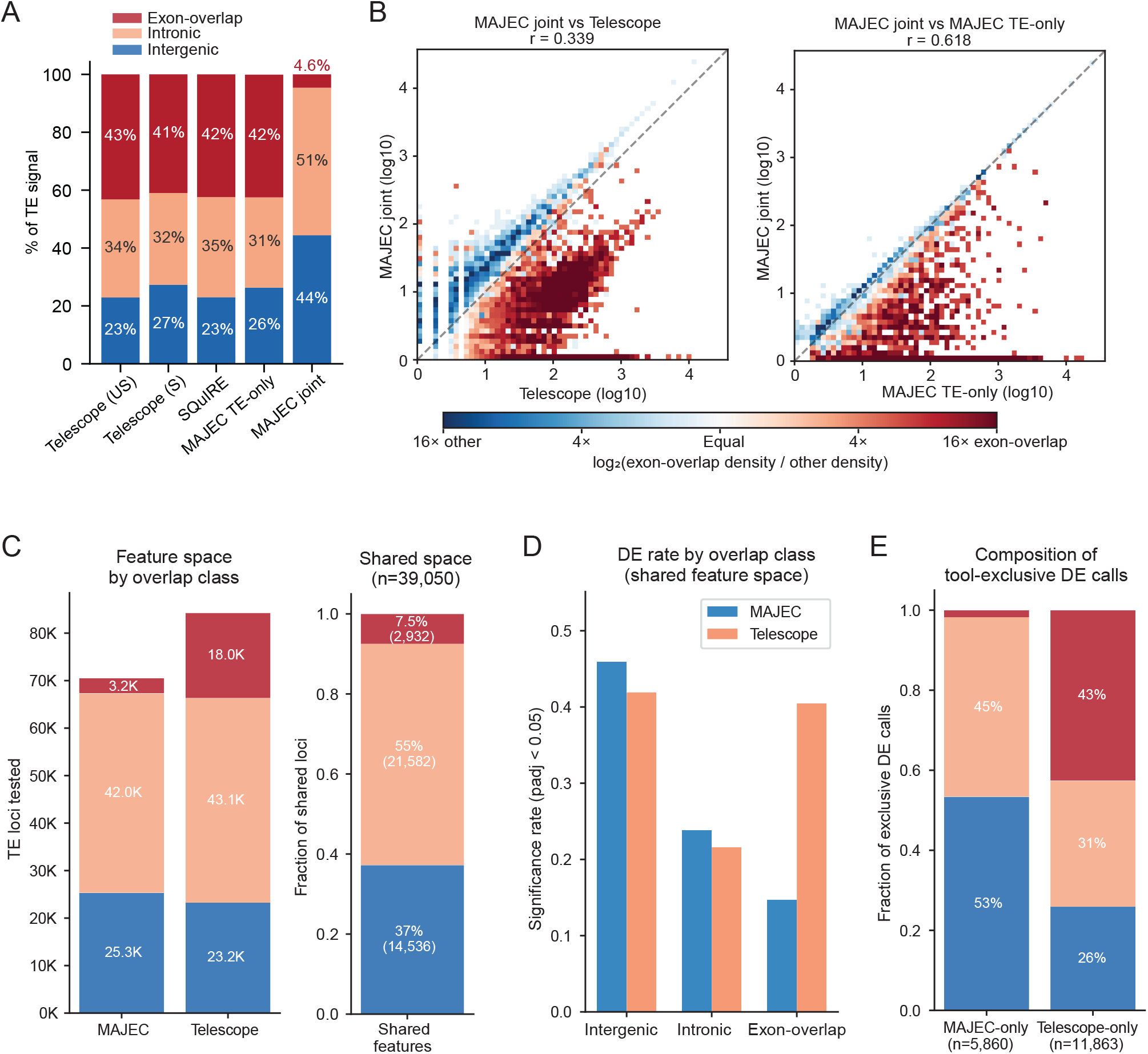
Joint gene-TE modeling eliminates exon-overlap contamination in locus-level TE quantification. **(A)** Fraction of total TE signal (counts) by genomic overlap class: intergenic (not within any annotated gene), intronic (within a gene body but not overlapping an annotated exon), or exon-overlapping (overlapping an annotated exon on either strand). Values are for five TE-quantification methods on a single library, with all loci classified against the same refGene gene model. Telescope (US) is unstranded; Telescope (S) is stranded (--stranded_mode RF, v1.0.3). The four TE-only methods attribute >40% of TE signal to exon-overlapping loci, including SQuIRE despite its use of a gene annotation; MAJEC joint reduces this to 4.6%. **(B)** Density scatter plots of per-locus counts between indicated tools, colored by local enrichment of exon-overlapping versus non-overlapping TE loci. Red indicates exon-overlap enrichment; blue indicates intergenic/intronic enrichment. Points below the diagonal represent exon-overlapping loci where Telescope or MAJEC TE-only assigns substantially more signal than MAJEC joint. **(C)** Left: total TE loci passing count thresholds for DESeq2 analysis, by overlap class. Right: overlap class composition of the 39,050 loci in the shared feature space. **(D)** Proportion of shared-space loci called significant (padj < 0.05) by each tool, stratified by overlap class. **(E)** Overlap class composition of DE calls exclusive to each tool (called significant by one tool but not the other, including tool-exclusive feature space).

Because Fig. 3A is based on real data, exon-overlapping TE signal could in principle include genuine TE transcription rather than contamination. To measure specificity directly, we simulated 40 million paired-end fragments from GENCODE coding and lincRNA transcripts only, with no TE sequence present, so any TE assignment is a false positive (Fig. S9). MAJEC misassigned 0.11% of fragments to TEs, compared with 8.0% for released (unstranded) Telescope and 4.2% for a development stranded build (75-fold and 40-fold higher misattribution). Strand-awareness roughly halved Telescope’s rate but did not change the composition of its errors, which were concentrated in ancient, gene-embedded Alu, MIR, and L2 elements. MAJEC’s smaller residual was instead dominated by young L1 subfamilies (L1HS and L1PA, 58%), likely reflecting cases where L1 sequence makes up a large enough fraction of the source transcripts that the reads cannot be confidently reassigned to a non-TE feature.

Exonic overlap was a major contributor to differences in expression estimates between the MAJEC joint model and Telescope or MAJEC TE-only mode (Fig 3B). The joint model’s effect is confined to exon-overlapping loci. Comparing MAJEC joint to MAJEC TE-only, intergenic and intronic loci are virtually unchanged (r = 0.99 and 0.96, median log2FC = 0.00 for both), while exon-overlapping loci are sharply reduced (median log2FC = −2.32; Fig. S10A). Against Telescope, the differences are larger across all classes (intergenic r = 0.52, intronic r = 0.39, exon-overlapping r = 0.34; Fig. S10B), reflecting additional algorithmic distinctions between the tools — strand awareness, EM priors, initial assignments — beyond the gene model itself. Intergenic signal increased under the joint model (464K to 639K fragments, median log2 ratio = +0.42), consistent with multi-mapper redistribution from deflated exonic copies to intergenic loci where no gene competes.

Exon-overlap contamination is therefore an architectural property of quantifying TEs outside a joint gene– TE feature space, shared by gene-unaware (Telescope), TE-only (MAJEC TE-only), and even gene-annotation-using (SQuIRE) methods alike.

The contamination flows through to differential expression analysis. We performed DESeq2 comparing the DMSO and decitabine treated cells on TE features having at least 5 counts in 2 or more samples. The number of features was generally similar between MAJEC (71K) and Telescope (84K) with Telescope having an enrichment in exon-overlapping TE consistent with the count analyses above (Fig. 3C). There were 39,050 TE loci shared between MAJEC and Telescope, and in that overlapping feature space the two methods detected similar overall numbers of significant DE loci (MAJEC: 12,256; Telescope: 11,948; padj < 0.05) but with substantially differing composition (Fig. 3C, D). Telescope called nearly three times more exon-overlapping loci as significant (1,187 vs. 432, or 40.5% vs. 14.7% of shared exon-overlap loci), while MAJEC detected more significant intergenic (6,678 vs. 6,095) and intronic (5,146 vs. 4,666) loci. Among the 1,872 loci called significant by Telescope but not MAJEC, 43% were exon-overlapping — a nearly 3-fold increase over the number called by MAJEC (Fig. 3D, S11A).

An additional 45,215 TE loci passed the count threshold in Telescope but not MAJEC; Telescope called 9,991 of these as significant, with exon-overlapping loci showing the highest rate (28.1% significant vs. 14.3% for intronic). In total, Telescope produced approximately 12,000 DE calls unsupported by MAJEC, with exon-overlapping loci disproportionately represented. Conversely, MAJEC detected 3,680 significant DE calls at 31,457 loci that fell below the count threshold in Telescope (Fig. S11B). These were overwhelmingly intergenic (58%) and intronic (41%), with only 254 exon-overlapping loci (0.8%) — consistent with multi-mapper redistribution directing signal toward intergenic and intronic loci where autonomous TE transcription is more plausible. In contrast to Telescope’s exclusive feature space, the DE calls private to MAJEC were relatively modest in significance, reflecting low-count loci that gained marginal signal through multi-mapper redistribution rather than large-effect reactivation events (Fig. S12).To illustrate the practical consequences at individual loci, we examined two vignettes from the same decitabine experiment that together demonstrate the complementary failure modes of existing tools (Fig. 4).

**Figure 4.**
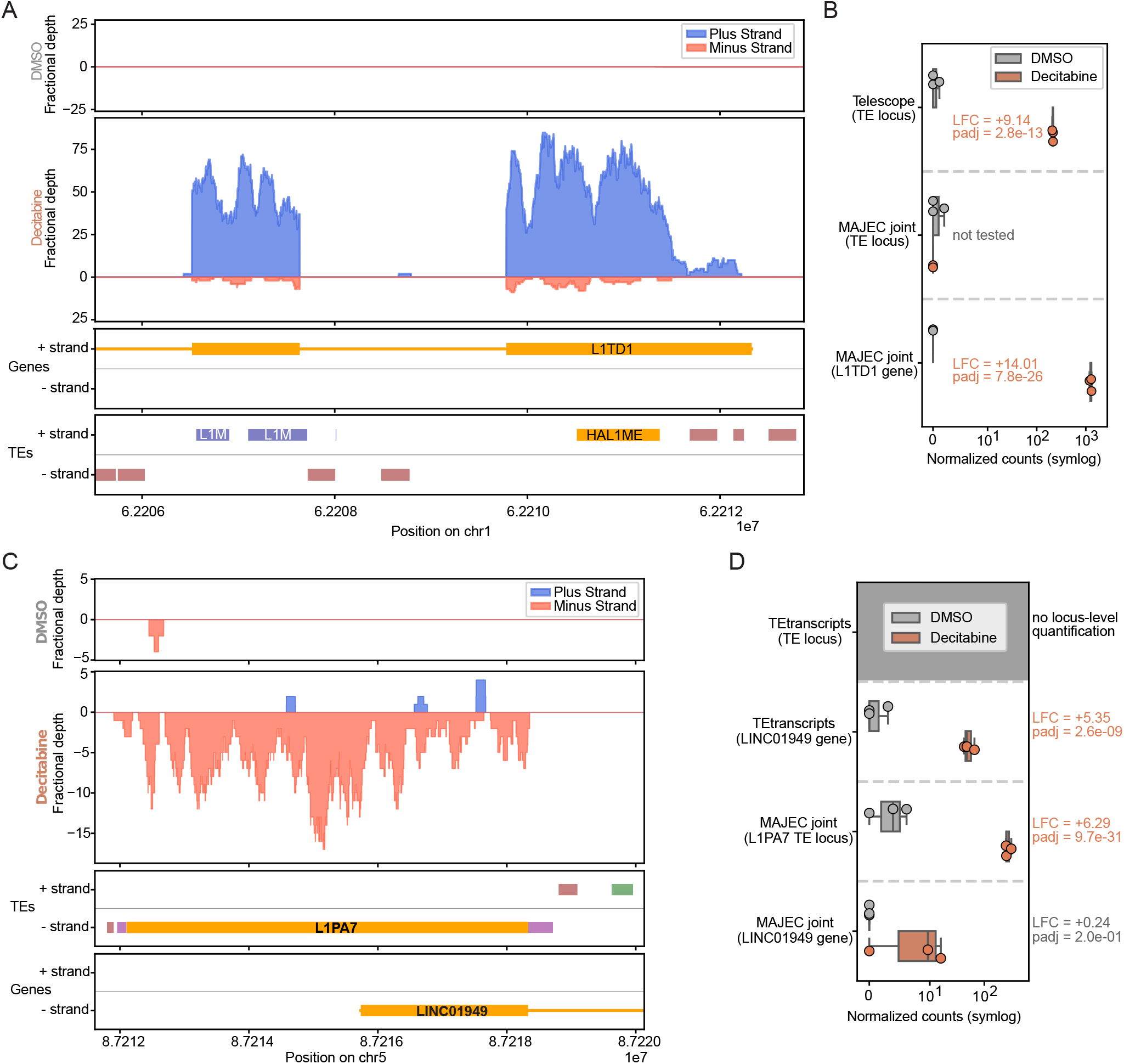
Complementary failure modes of gene-unaware and rule-based TE quantification. (A) Stranded fractional read coverage at the 3’ end of L1TD1 (chr1) in DMSO (top) and decitabine (middle) conditions, with gene and TE annotation tracks below. Plus-strand coverage in decitabine shows a clear exon-intron pattern matching L1TD1; no independent signal is visible over the HAL1ME or L1MEd annotations. (B) Normalized counts (symlog scale) for the HAL1ME locus from Telescope and MAJEC, and for the L1TD1 gene from MAJEC. Telescope reports HAL1ME as highly significantly upregulated (LFC = +9.14, padj = 2.8×10⁻¹³); MAJEC assigns insufficient counts to test the TE locus and instead reports massive upregulation of L1TD1 (LFC = +14.01, padj = 7.8×10⁻²⁶). (C) Stranded fractional read coverage at LINC01949 (chr5), showing diffuse minus-strand coverage spanning the L1PA7 element in decitabine without splice structure matching the LINC01949 annotation. (D) Normalized counts for the L1PA7 locus and LINC01949 gene from TEtranscripts and MAJEC. TEtranscripts lacks locus-level TE resolution (gray, top row) and misattributes L1PA7-derived signal to LINC01949 (LFC = +5.35, padj = 2.6×10⁻⁹). MAJEC assigns signal primarily to the L1PA7 locus (LFC = +6.29, padj = 9.7×10⁻³¹) and reports LINC01949 as unchanged (LFC = +0.24, padj = 0.20).

### Vignette 1: False TE reactivation (Telescope’s failure)

HAL1ME_dup104 is an element residing within the second exon of L1TD1, a pluripotency-associated gene silenced by DNA methylation in most somatic cells (Altenberger *et al*. 2017, Kavaklioglu *et al*. 2025). Telescope reports this TE locus as highly significantly upregulated by decitabine treatment (LFC = +9.1, adjusted p = 2.8 × 10⁻¹³; Fig. 4A–B). In isolation, this suggests the demethylation-induced reactivation of an ancient LINE element. However, MAJEC’s joint model correctly assigns these reads to L1TD1, which shows massive upregulation (LFC = +14.0, adjusted p = 7.8 × 10⁻²⁶). Because L1TD1’s transcript model has strong junction support across all decitabine samples, it receives a pre-EM junction boost, allowing MAJEC to recognize the signal as unambiguously genic. Stranded coverage confirms this: decitabine-treated samples show strong plus-strand coverage with a clear exon-intron pattern matching L1TD1, with no isolated peak over the TE annotation (Fig. 4A–B). Similar patterns were consistently observed in exon overlapping TE of other decitabine derepressed genes such as IL11 (Fig. S13).

### Vignette 2: False gene upregulation (TEtranscripts’ failure)

The converse error occurs when a genuinely reactivated TE is embedded within a gene. L1PA7_dup4347 is a minus-strand L1PA7 element residing within LINC01949, a noncoding RNA. MAJEC’s joint model correctly separated the two signals: the TE locus was strongly upregulated (LFC = +6.29, padj = 9.7 × 10⁻³¹), while LINC01949 showed no change (LFC = +0.24, padj = 0.20; Fig. 4C–D). Notably, LINC01949 has two annotated splice junctions but zero junction-spanning reads in any sample, triggering a completeness penalty that reduced its prior — the same junction system that assigns reads to L1TD1 in vignette 1 withholds them from LINC01949. Stranded coverage confirmed genuine TE transcription: minus-strand signal in decitabine samples spanned the L1PA7 annotation without exon-intron structure, consistent with cryptic transcription from a demethylated TE promoter rather than spliced lincRNA expression (Fig. 4C). Because TEtranscripts assigns uniquely mapped reads at gene-TE overlaps exclusively to the gene, these TE-derived reads were absorbed into the LINC01949 count, producing a false DE call (LFC = +5.35, padj = 2.6 × 10⁻⁹). A researcher using TEtranscripts would conclude they had discovered a decitabine-responsive lincRNA; the real biology was an embedded L1 element being demethylated.

These two vignettes illustrate opposing failure modes that MAJEC’s joint model resolves simultaneously. Telescope, operating without gene annotations, misattributes genic reads to overlapping TEs, while TEtranscripts, applying a rigid gene-over-TE prioritization rule, absorbs genuine TE signal into host genes. The Telescope-direction error is pervasive affecting multiple TE per upregulated gene (Fig 4A, S10). The TEtranscripts-direction error is much less common but systematically biased. Of 52 loci where MAJEC identified TE-dominated signal at a genic locus, TEtranscripts reported significant gene upregulation of the overlapping host gene at 18 (Table S1). Among the eight host genes with the largest inflated log-fold-change calls, seven were lincRNAs, and the implicated TE families (LTR1A2, LTR12C, HERVH-int, L1PA7, L1PB1, HAL1) include several with documented reactivation or cryptic-promoter activity upon DNA demethylation (e.g., LTR12C; Brocks et al. 2017; Chuong et al. 2017). (Table S1).

MAJEC resolves both cases correctly because it does not impose a fixed rule about which feature type prevails. Instead, genes and TE loci compete for reads probabilistically within the joint EM, and the assignment follows the evidence. When the evidence supports genic origin (L1TD1: exon-intron structure, strand-concordant coverage), reads flow to the gene. When the evidence supports TE origin (LINC01949/L1PA7: localized TE-region coverage without splice structure, silent host gene), reads remain with the TE. A probabilistic model gets both cases right because it asks “what does the data support?” rather than applying a predetermined answer.

### Runtime and computational requirements

MAJEC provides practical computational efficiency alongside its accuracy improvements. We compared the peak memory usage and runtimes from MAJEC, TEtranscripts, and Telescope for our decitabine dataset (Table 1). Both TEtranscripts and Telescope process samples individually on single cores, but their computational bottlenecks differ significantly. Telescope is trivially parallelizable across samples, requiring approximately 31 minutes to process a single BAM. However, its ∼13 GB per-sample memory footprint means that running six samples simultaneously demands ∼78 GB of RAM, effectively exceeding the capacity of a standard 64 GB workstation. Conversely, TEtranscripts is highly memory efficient (∼6 GB per sample), likely owing to its collapse of the TE feature space to the subfamily level. Yet, because it is inherently designed for differential expression, it requires at least one test and one control BAM and processes them serially at roughly one hour per sample. This dictates a minimum runtime of two hours even in a parallelized workflow.

**Table 1.**
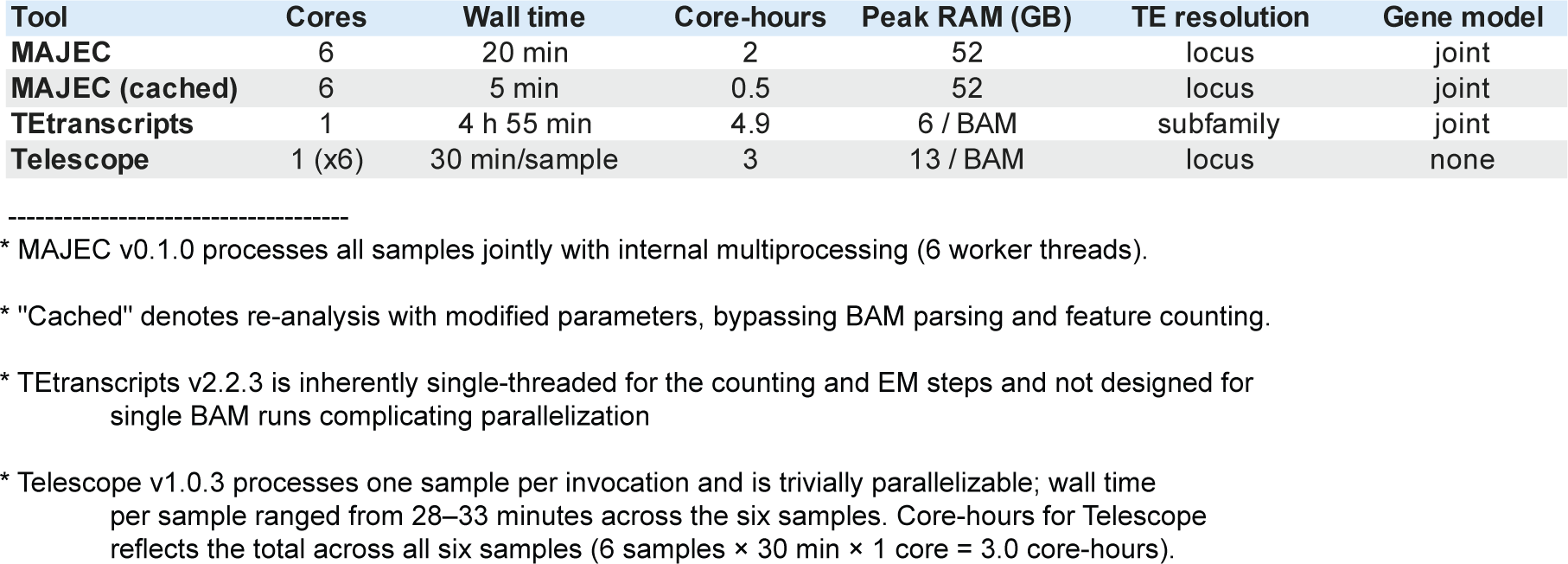
Computational performance and feature comparison of TE quantification tools. Runtime, memory, and model characteristics for processing six BAM files on an Intel Xeon HPC cluster. MAJEC utilizes internal multiprocessing to achieve rapid locus-level, joint gene-TE quantification (20 min wall time, 2.0 core-hours). Its caching mechanism further reduces re-analysis time to 5 minutes by bypassing initial BAM parsing. TEtranscripts relies on single-threaded execution, requiring 4 hours 55 minutes for subfamily-level resolution. Telescope processes samples individually (∼30 minutes/sample) but operates in a TE-only feature space. Peak RAM is reported for the joint run (MAJEC) or per individual BAM file (TEtranscripts, Telescope).

In stark contrast, MAJEC overcomes both of these limitations. Its multithreaded architecture processes all six samples jointly in just 20 minutes across 6 cores, utilizing a single, shared memory footprint of ∼50 GB. MAJEC’s efficiency advantage is thus roughly 4× in core-hours compared to the combination of Telescope and TEtranscripts needed to approximate its output—while also providing additional wall-time benefits and isoform-level quantification that neither competing tool offers.

Runtime is dominated by I/O rather than computation (Table S2). The momentum-accelerated EM completed in less than 1.5 minutes across 15–16 iterations, accounting for less than 10% of wall time. Peak memory (∼50 GB) is within the range of a standard STAR alignment; MAJEC does not require HPC infrastructure. A caching mode that stores featureCounts assignments and junction metrics enables rapid parameter exploration (∼5 minutes per rerun vs ∼20 minutes for a full run).

## 4. Discussion

MAJEC demonstrates that gene, isoform, and locus-level TE quantification can be unified in a single analysis that is faster, more comprehensive, and more accurate at gene-TE boundaries than running specialized tools in sequence. Three aspects of these results warrant further discussion: the mechanistic basis of MAJEC’s isoform accuracy, the scope and implications of the gene-TE overlap problem, and the practical limitations of the current implementation.

### Unified quantification is faster and simpler

The most immediate practical benefit of MAJEC is consolidation. A typical RNA-seq workflow studying TE biology requires STAR alignment (for visualization and QC), Salmon (for isoform quantification), TEtranscripts (for subfamily-level TE DE), and potentially Telescope (for locus-level TE resolution) — four tools with three different annotation formats and incompatible output structures. MAJEC replaces the three downstream tools with a single analysis that operates on the existing BAM. At comparable CPU allocation to TEtranscripts (1 core / BAM), MAJEC completes in approximately 20% of the time, without sacrificing subfamily-level concordance with TEtranscripts (r > 0.99) or locus-level resolution.

Beyond quantification, MAJEC provides integrated downstream tools: a queryable SQLite database storing all count estimates, junction evidence, and confidence metrics; automated extraction of DESeq2-ready count matrices at gene, isoform, and TE locus levels; and interactive HTML reports for isoform-level QC and exploratory analysis. For users whose primary interest is isoform resolution rather than TE biology, MAJEC and Salmon are complementary: Salmon’s sequence-level modeling and GC bias correction capture information that MAJEC’s alignment-based approach does not, while MAJEC provides the joint gene+TE model that alignment-free tools structurally cannot. Running both from the same BAM adds only minutes to the analysis.

### Junction penalties are the core innovation — and the mechanism is clear

MAJEC’s isoform accuracy advantage over Salmon and RSEM is not a general improvement across all transcripts but a targeted effect confined to specific transcript classes. On simple synthetic transcriptomes (Sequins, max 4 isoforms per gene), all methods performed equivalently. On complex real transcriptomes (LongBench, 8 ENCODE cell lines), MAJEC outperformed Salmon on 54% of transcripts — but this advantage was entirely driven by transcripts receiving incomplete or subset junction penalties. Transcripts with no junction penalty showed no MAJEC advantage whatsoever. This stratification establishes that the junction prior system helps precisely where isoform structure is ambiguous and is inert where it is not.

Feature ablation confirmed this interpretation from the causal direction. Removing completeness penalties significantly degraded MAJEC accuracy resulting in performance that was inferior to Salmon by approximately 10% while disabling the subset penalties caused further erosion. The innovation is not better evidence for well-supported transcripts; it is better use of evidence to downweigh poorly-supported ones.

The penalty-type stratification and ablation results replicate across two independent datasets (T-cell and LongBench), and the advantage is consistent across all four long-read ground truths (dRNA, PacBio, ONT, and their consensus), with dRNA — arguably the least biased long-read platform due to the absence of reverse transcription and PCR amplification (Garalde *et al*. 2018, Workman *et al*. 2019) — showing the largest MAJEC advantage.

The precision-sensitivity tradeoff inherent in the penalty system is an intentional design choice. By aggressively penalizing unsupported isoforms, MAJEC calls 7,000–9,000 fewer false-positive transcripts per cell line than Salmon. This precision is essential for the joint gene+TE model, as it prevents spurious genic transcripts from inappropriately absorbing reads that belong to legitimate TE loci. However, this comes at the cost of suppressing some genuinely expressed, low-abundance transcripts. Users focused purely on discovering rare, novel isoforms may prefer Salmon’s higher sensitivity, while those analyzing TE biology will benefit from MAJEC’s strict genic boundaries. Implementing an expression-dependent penalty floor to reduce penalty strength when total evidence is sparse represents a natural future improvement.

### The gene-TE overlap problem is larger than appreciated

The most striking finding of this study is the sheer scale of exonic overlap contamination in locus-level TE quantification. When using Telescope, exon-overlapping loci—which constitute only 1.1% of annotated TE loci—account for 43% of the total TE signal. MAJEC’s joint model reduces this exonic TE signal to just 5% and successfully relocates multi-mapped reads from gene bodies to intergenic loci where autonomous TE transcription is biologically plausible. Because these fractions come from cellular RNA, where a portion of the exon-overlapping signal could in principle be genuine, we turned to a ground-truth control in which RNA-seq was simulated from gene models with no TE transcribed. In that setting MAJEC attributed almost none of the gene-derived signal to TEs (0.11%), nearly two orders of magnitude below Telescope, suggesting that most exon-overlapping TE signal reflects contamination rather than authentic TE expression (Fig. S9).

This contamination severely confounds differential expression analysis. Despite testing in the same DESeq2 framework, Telescope called nearly 12 times more exon-overlapping loci as significantly differentially expressed compared to MAJEC (5,415 vs. 461). This massive exonic inflation accounts for over 80% of the total discrepancy in DE calls between the two tools. This contamination acts as a double-edged sword: inflated exonic counts artificially generate false positives while simultaneously attracting multi-mapped reads away from intergenic loci during the EM, reducing the statistical power to detect genuine TE activation.

Genic contamination is not a Telescope-specific artifact but a general property of TE-only quantification. We observed the same effect with SQuIRE, which attributes >40% of its TE signal to the exon-overlapping class despite being gene-aware. While SQuIRE ingests a gene annotation, it is only used to quantify genes and to guide splice-aware alignment not to allow arbitration of reads among overlapping gene and TE annotations. Telescope, likewise, was designed to solve the multi-mapper reassignment problem, and does so effectively through its Bayesian model, but that model resolves ambiguity among competing TE loci, not between TEs and overlapping genes. Neither tool performs gene-versus-TE read arbitration and the scale of this contamination was not previously quantified. Our results suggest that published locus-level TE analyses using any TE-only tool should be interpreted with caution at gene-overlapping loci, and that future studies would benefit from either joint gene-TE modeling or, at minimum, post hoc filtering of exon-overlapping TE signal.

The overlap problem is also bidirectional. While Telescope and SQuIRE inflate TE signal at loci where genes are expressed, TEtranscripts’ gene-over-TE prioritization rule can absorb genuine TE transcription into host gene counts, creating spurious gene-level differential expression calls. We identified several such cases among 52 loci where MAJEC detected TE-dominated signal at genic loci, where 7 of the 8 most affected host genes were lincRNAs. This systematic bias toward noncoding genes is expected: the majority of lincRNA exonic sequence is TE-derived, making lincRNAs uniquely vulnerable to a rule that unconditionally prioritizes gene annotations (Kelley and Rinn 2012). Several of the affected TE families — notably LTR12C and the LTR1/HERV LTRs — have documented promoter or cryptic-transcription activity that is derepressed upon DNA demethylation (Chuong *et al*. 2017). These are precisely the TEs most likely to be genuinely reactivated by decitabine and, because much lincRNA sequence is TE-derived (Kelley and Rinn 2012), most likely to reside within the noncoding genes susceptible to this misattribution. Studies of TE reactivation in contexts where noncoding transcriptome analysis matters — epigenetic therapy, cancer, development — should be particularly attentive to this class of artifact (Brocks *et al*. 2017).

The convergence between MAJEC and TEtranscripts at the subfamily level, despite fundamentally different approaches to genic contamination, provides strong validation that the joint model produces correct TE estimates. Meanwhile, Telescope diverges from both (r ≈ 0.92), confirming that the grouping variable is not the algorithm but whether gene-TE overlap is addressed at all.

### Limitations and future directions

Several limitations of the current implementation suggest directions for improvement. The EM can over-concentrate counts for asymmetric isoform pairs where one transcript has substantially more unique evidence than the other, iteratively driving the minor isoform to zero. This affects a small number of transcripts (approximately 1,000 across 8 cell lines, identifiable through the discord score) and could be addressed with pseudocount regularization. More fundamentally, the current architecture separates feature assignment from junction evidence: featureCounts assigns reads to equivalence classes based solely on exon overlap, discarding the splice junction information encoded in read CIGAR strings and the per-base alignment quality (mismatch counts and alignment scores) that could distinguish near-identical paralogs and processed pseudogenes from their parent loci before the EM stage. A read spanning two exons connected by a transcript-diagnostic junction is indistinguishable from one overlapping the same exons without splicing. For paired-end libraries, intersecting the transcript sets from both mates of a fragment provides additional narrowing, as a fragment can only originate from a transcript compatible with both ends. A prototype single-pass BAM tool implementing this approach consolidates the current three-pass pipeline (unique mapper assignment, multimapper filtering, multimapper assignment) into a single scan that simultaneously resolves exon overlap, junction compatibility, and read-pair agreement. Incorporating alignment metrics into equivalence-class weighting through a direct BAM parser is a natural extension of the single-pass prototype noted above. Whether tighter equivalence class resolution or incorporation of alignment quality can translate to improved quantification accuracy remains to be validated against long-read ground truth. Parameter sensitivity analysis also identified marginal improvements on the LongBench dataset, though these may be dataset-specific and require cross-validation before changing defaults.

MAJEC is alignment-based and therefore slower than Salmon for pure isoform quantification. However, TE quantification inherently requires genomic alignment — multi-mapped reads to repetitive genomic loci cannot be resolved by alignment-free approaches that operate on transcript-level indices — so the comparison for users needing TE analysis is not MAJEC versus Salmon but MAJEC versus the combined multi-tool pipeline. In this comparison, MAJEC is several times faster.

The joint model’s effectiveness depends on the quality of both gene and TE annotations. On well-annotated genomes (human, mouse) with comprehensive GENCODE transcript models and RepeatMasker TE libraries, the model can confidently assign reads at gene-TE boundaries. For non-model organisms with incomplete gene annotations, the benefit of joint modeling is reduced — poorly annotated genes cannot absorb their genic TE reads. Comparisons between MAJEC and Telescope output were performed without strand discrimination, as Telescope lacks strand awareness; this introduces ambiguity at loci where TEs overlap antisense genes and may account for some disagreements between the two methods.

Finally, long-read sequencing, which increasingly resolves multi-mapping ambiguity through read length alone, represents a complementary approach to the same problems MAJEC addresses computationally; recent tools such as LocusMasterTE (Lee *et al*. 2025) leverage long-read data for improved locus-level resolution. TEspeX (Ansaloni *et al*. 2022) addresses the exonized TE fragment problem through a filtering rather than modeling approach. Both tools recognize the gene-TE overlap problem from different angles, and MAJEC’s probabilistic joint modeling represents a third strategy that is complementary to both.

## Acknowledgments

We thank David Hendrickson for helpful discussions and manuscript review. The authors acknowledge the use of generative AI technologies, specifically Claude (Anthropic) and Gemini (Google), for assistance with code generation, manuscript drafting, and the exploration of analytical approaches.

---

**Figure S1.**
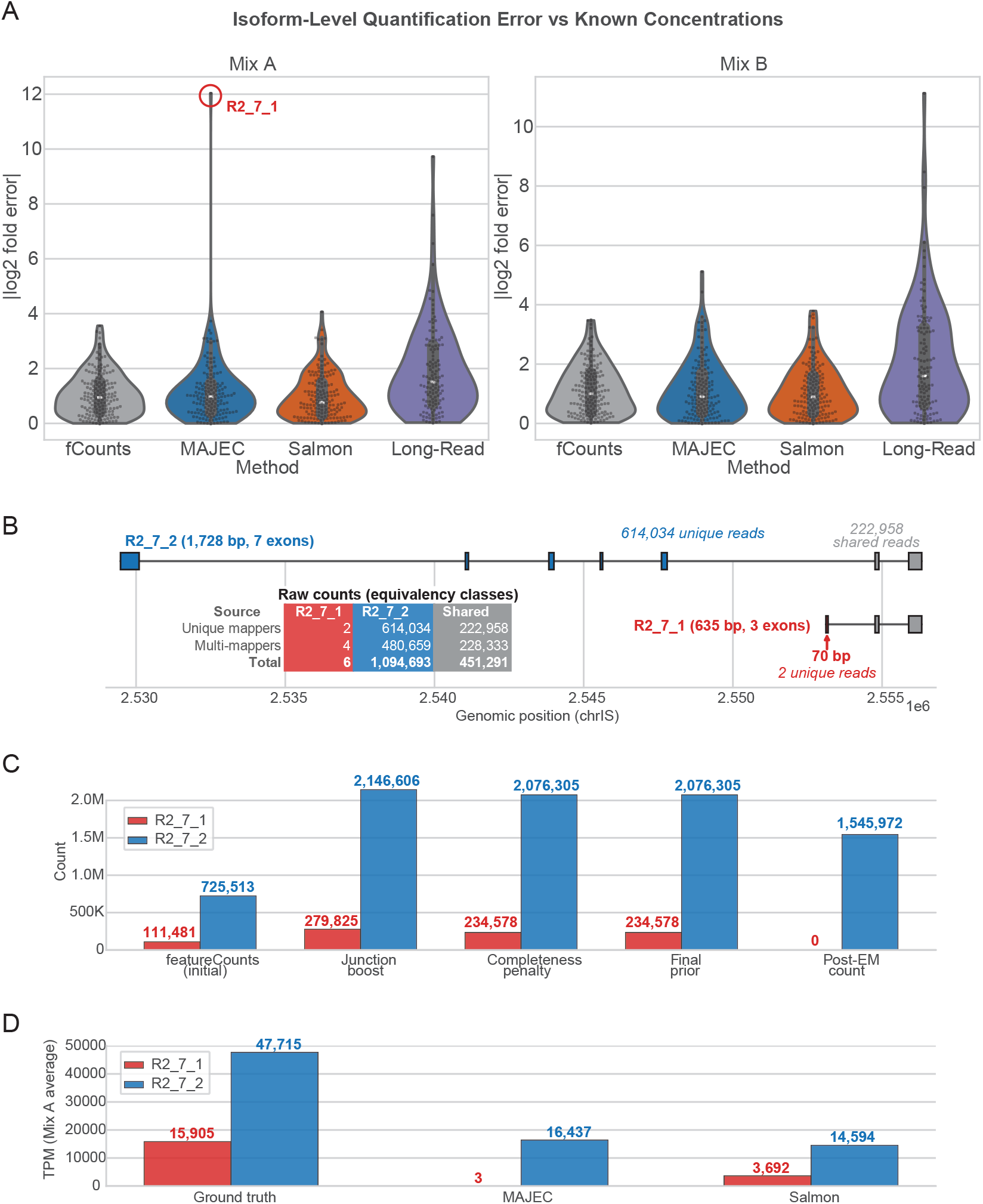
EM over-concentration at asymmetric isoform pairs. **(A)** Violin plots of per-isoform absolute log2 fold error for each quantification method on the Sequins v2.4 dataset (Mix A and Mix B). The R2_7_1 outlier (circled, Mix A) represents MAJEC’s worst-case failure. **(B)** Genomic structure of the R2_7 locus (chr18), showing two isoforms: R2_7_2 (7 exons, 1,728 bp) with 614,034 unique reads and 222,058 shared reads, and R2_7_1 (3 exons, 635 bp) with only 70 bp of unique territory and 2 unique reads. **(C)** Per-isoform count progression through MAJEC’s prior adjustment stages. R2_7_2 receives large junction boosts from its exclusive junctions; R2_7_1 receives only a modest boost. The completeness penalty applies minor suppression to R2_7_1 (0.84), and the EM drives its final count to zero. **(D)** Final TPM estimates compared to ground truth (Mix A). MAJEC assigns nearly all signal to R2_7_2 (47,715 TPM vs ground truth 16,906) and zeroes R2_7_1 (3 TPM vs ground truth 16,906). Salmon distributes counts more evenly but still favors R2_7_2.

**Figure S2.**
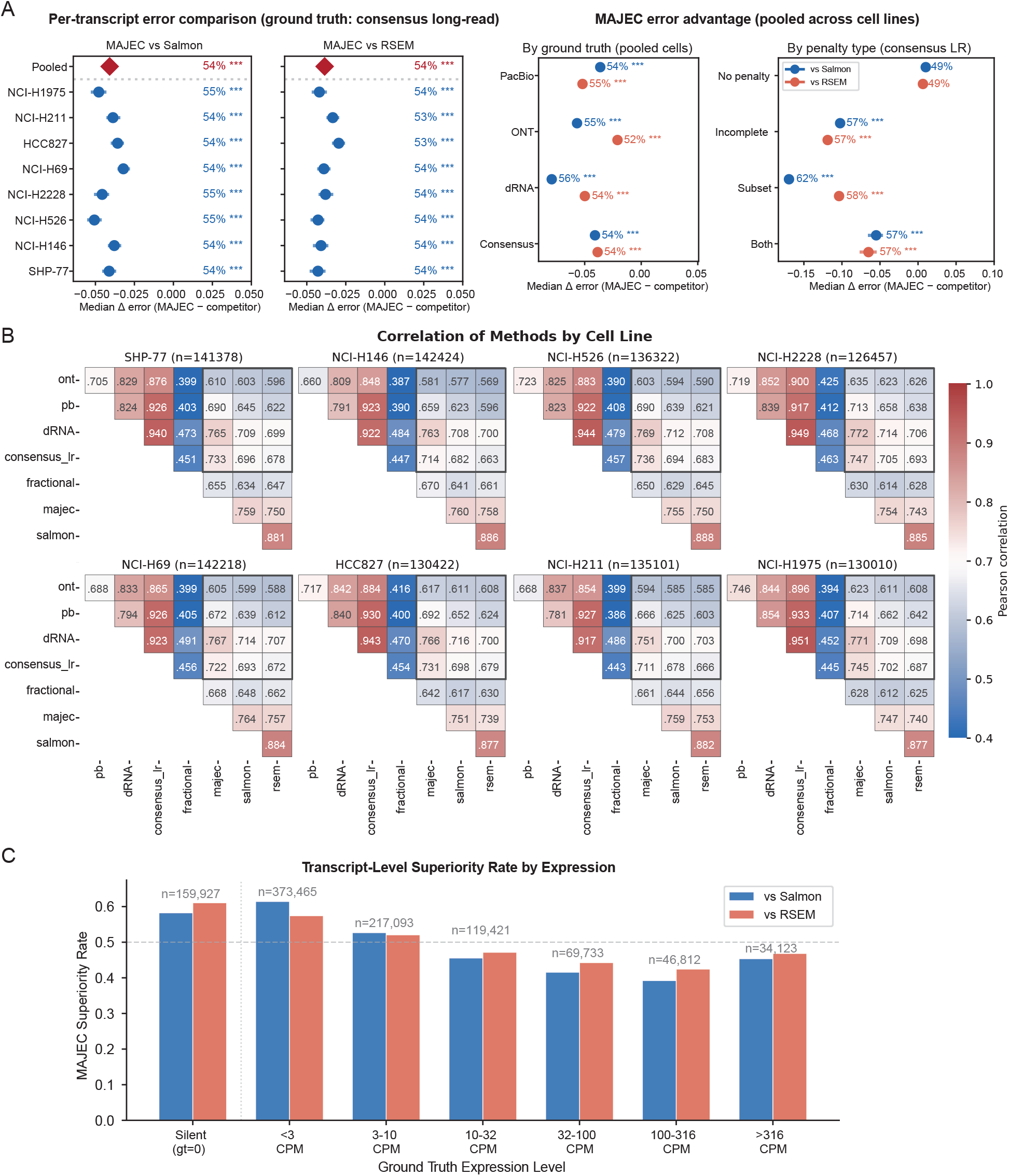
LongBench isoform quantification benchmarking: per-cell-line and per-platform details. **(A)** Forest plots of MAJEC’s median error advantage over Salmon and RSEM (Left) for each cell line individually and pooled, with 95% bootstrap confidence intervals (consensus long-read ground truth). Superiority rates and Wilcoxon signed-rank significance (*** padj < 0.001) are shown. Right: Superiority rates pooled for all cell lines and stratified by long-read platform and by MAJEC junction penalty type. MAJEC’s advantage is largest against dRNA ground truth and at subset-penalized transcripts; no significant advantage is observed for unpenalized transcripts. **(B)** Pairwise Pearson correlation heatmaps (log1p-transformed counts-per-10-million) for each cell line across all quantification methods and long-read platforms. **(C)** MAJEC superiority rate stratified by ground truth expression level. MAJEC’s advantage is largest at silent and low-expression transcripts and falls below 50% at mid-expression levels.

**Figure S3.**
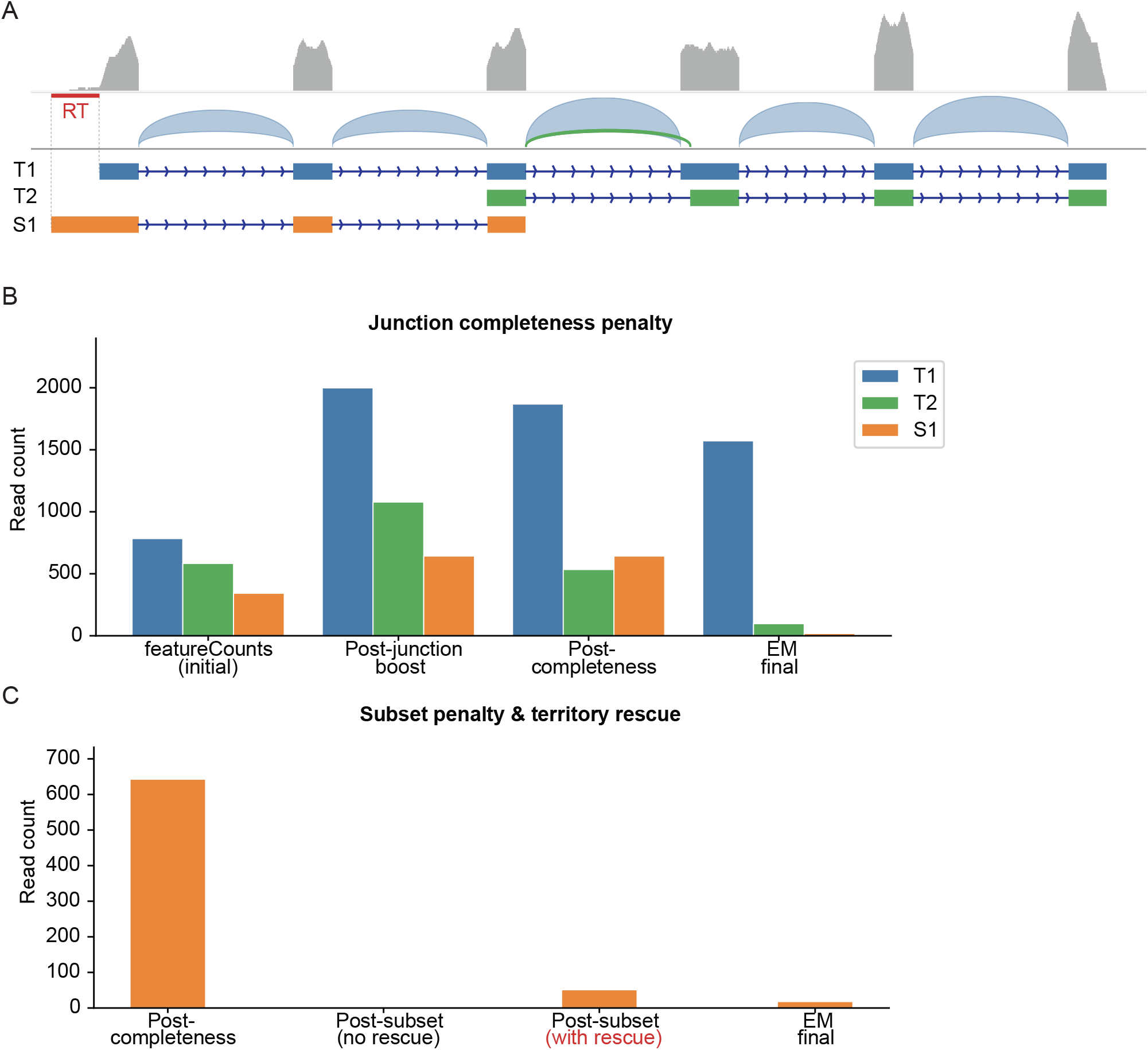
Demonstration of MAJEC prior adjustment on synthetic data. **(A)** IGV visualization of a synthetic locus containing three annotated transcripts of a synthetic gene: T1 (full-length, 6 exons), T2 (4 exons sharing E3 with T1 but using an alternative splice acceptor at E4’), and S1 (3 exons, a junction subset of T1 with a unique 5’ extension). Coverage track (gray) and splice junction arcs (blue/green) are shown above transcript models. The red region (RT) marks S1’s unique territory used for subset rescue. The green arc highlights T2’s weakly supported junction versus T1’s strongly supported E3→E4 junction at the same donor site. **(B)** MAJEC prior adjustment stages for junction completeness. Initial counts reflect featureCounts fractional assignment. Junction evidence boosts priors proportionally to splice junction support. The completeness penalty then downweights transcripts with poorly supported junctions relative to their gene context. **(C)** Subset penalty and territory rescue for S1. Because all S1’s splice junctions are contained within T1 and show limited evidence of intendent S1 contribution, S1 receives a near-complete subset penalty. However, read coverage in S1’s unique 5’ territory (red region in A) partially rescues the prior, reflecting evidence that S1 may represent a real isoform despite its junction redundancy. Final EM counts: T1 = 1,572, T2 = 98, S1 = 18

**Figure S4.**
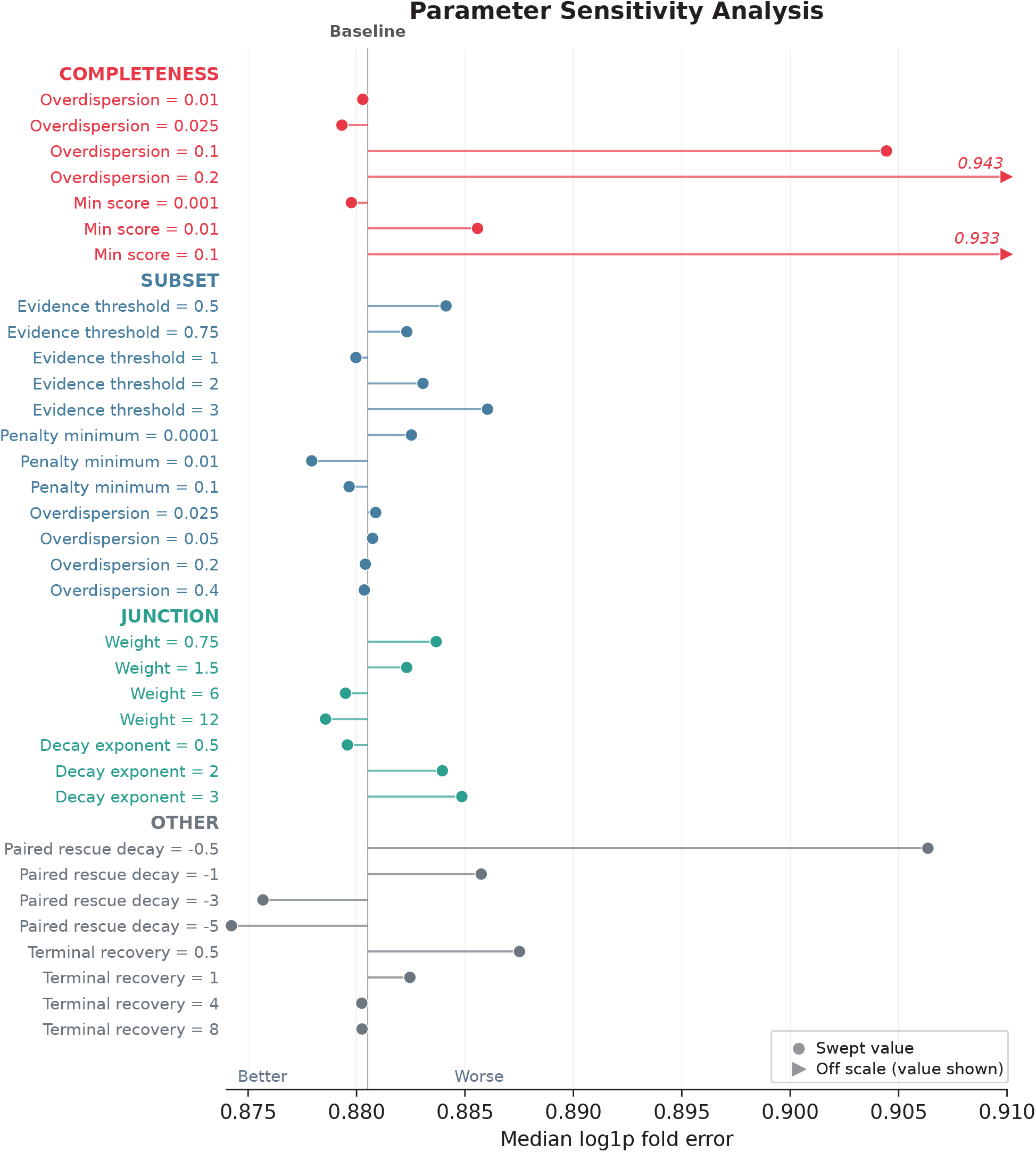
Parameter sensitivity analysis. Each point shows the median log1p fold error across 8 cell lines against the consensus of long read data for a single parameter setting, with all other parameters held at their default values. The vertical gray line marks the baseline configuration. Parameters are grouped by feature: completeness penalty (overdispersion, min score), subset penalty (evidence threshold, penalty minimum, overdispersion), junction boost (weight, decay exponent), and other (paired rescue decay, terminal recovery rate). Default values: completeness overdispersion = 0.05, completeness min score = 0.0001, subset evidence threshold = 1.25, subset penalty minimum = 0.001, subset overdispersion = 0.1, junction weight = 3.0, junction decay exponent = 1.0, paired rescue decay = −1.5, terminal recovery rate = 2.0. Circles indicate swept values; triangles indicate conditions whose error falls beyond the plotted axis. Performance is broadly robust to parameter perturbation, with the exception of completeness overdispersion ≥ 0.1 and completeness min score ≥ 0.1, which substantially degrade accuracy by under-penalizing incomplete transcripts.

**Fig. S5.**
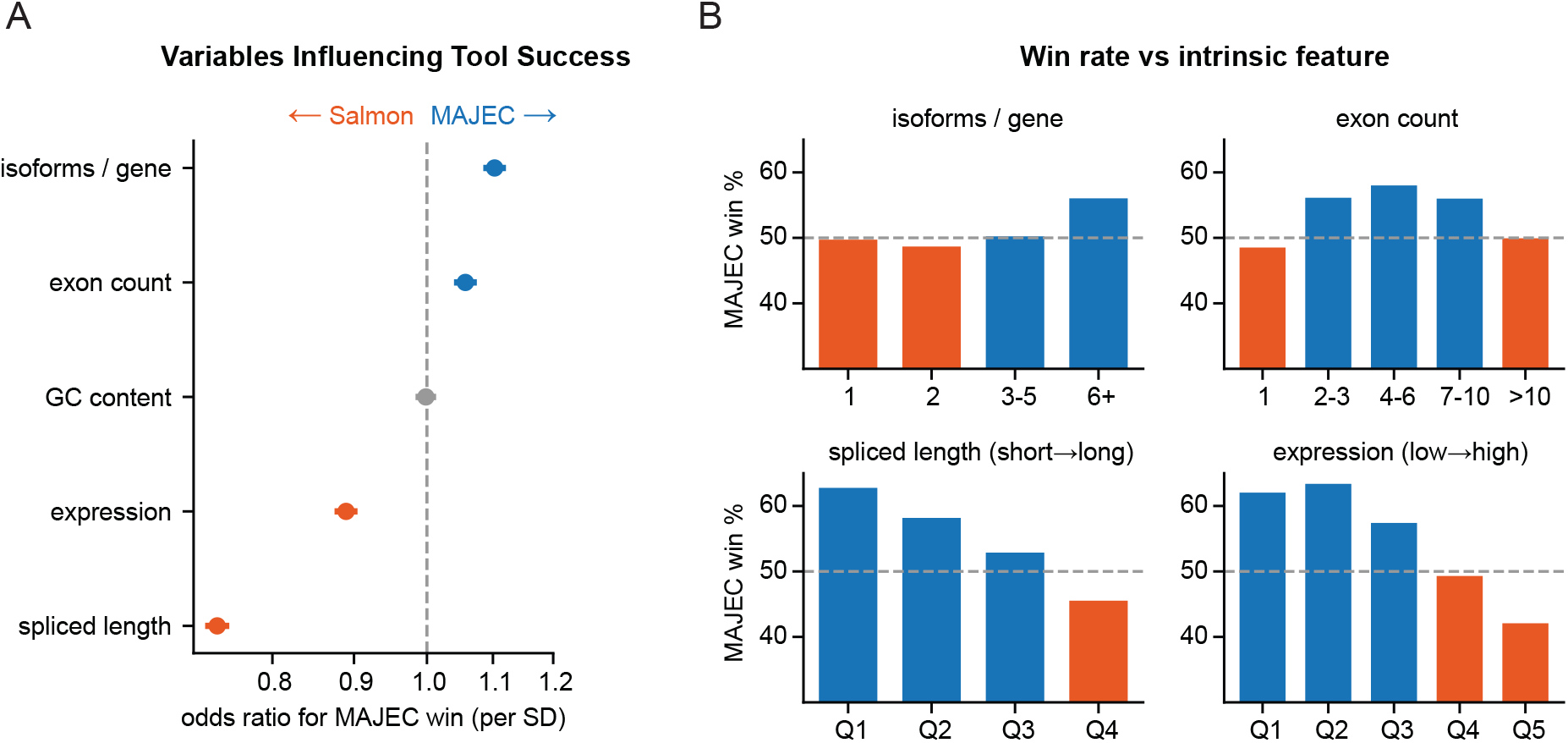
Intrinsic transcript features associated with MAJEC versus Salmon accuracy in LongBench. For each transcript in each cell line, the tool with the lower absolute error (a per-transcript “win”) was modeled against five standardized transcript features. The analysis is restricted to protein-coding and lncRNA transcripts, the biotypes reliably measured by long-read sequencing (n = 969,837 transcript-sample observations across 175,903 transcripts; overall MAJEC win rate 54.8%). **(A)** Odds ratios from a multivariable logistic regression for a MAJEC win, with z-scored predictors so each odds ratio is the change in odds of a MAJEC win per one standard-deviation (SD) increase in that feature. Odds ratios above 1 favor MAJEC (blue), below 1 favor Salmon (orange), and grey marks a 95% confidence interval spanning 1 (not significant); the dashed line is OR = 1. Standard errors are cluster-robust by gene (30,869 clusters) to account for transcripts observed across multiple cell lines, and confidence intervals are narrow given the large n. **(B)** Proportional MAJEC win rate (ties counted as 0.5) by isoform count per gene, exon count, spliced-length quartile, and expression quintile, with bars colored by the winning tool and the dashed line marking 50% parity.

**Figure. S6.**
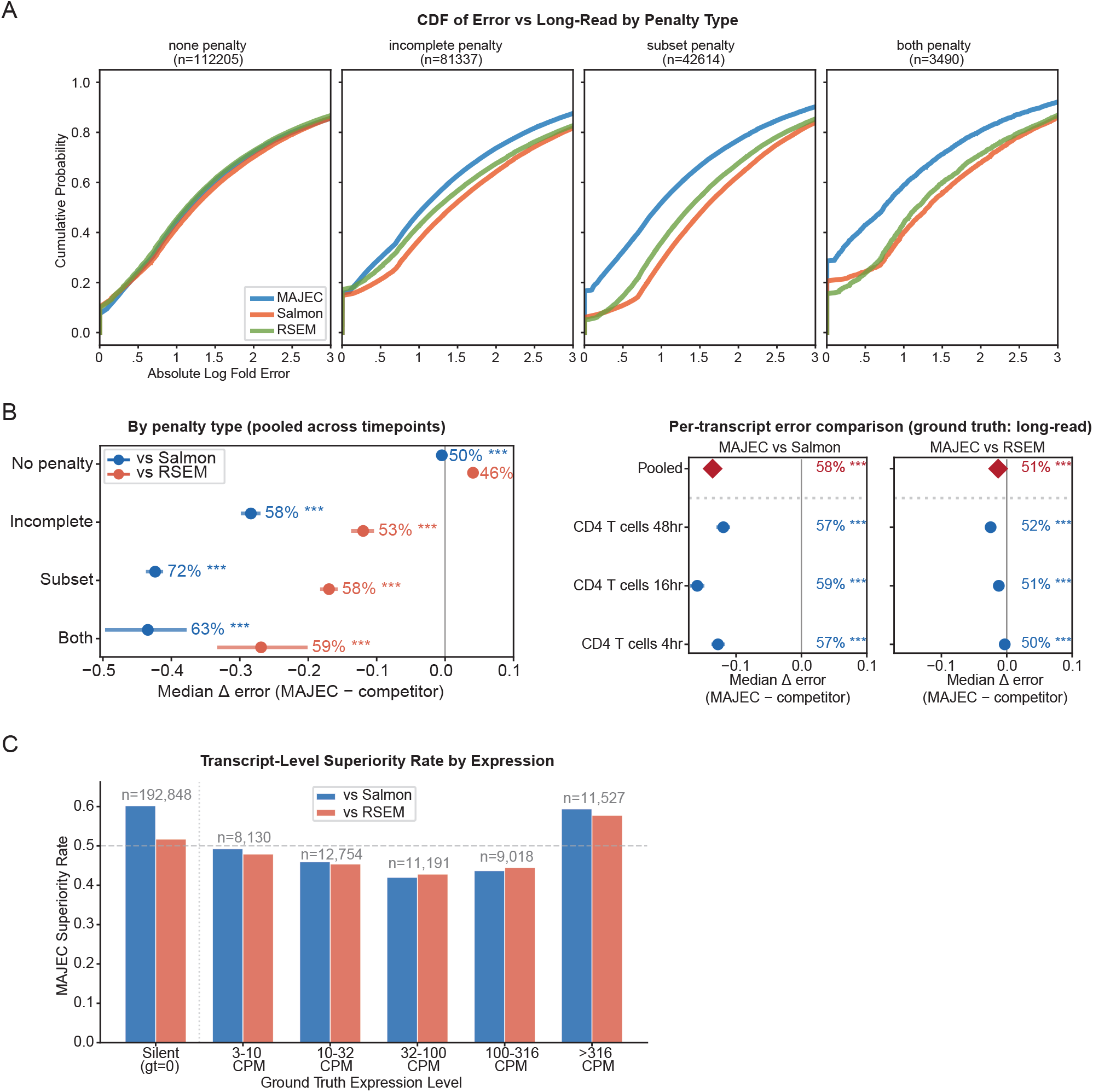
Benchmarking transcript-level quantification accuracy of MAJEC, Salmon, and RSEM using long-read ground truth. Transcript quantification from short reads was compared against matched PacBio long-read data (quantified via IsoQuant) across three CD4 T cell activation timepoints (4hr, 16hr, and 48hr). **(A)** Cumulative distribution function (CDF) of absolute log-fold error for MAJEC (blue), Salmon (orange), and RSEM (green) relative to long-read counts. Transcripts are stratified by MAJEC’s algorithmically assigned prior penalty type (none, incomplete, subset, or both). **(B)** Paired comparisons of per-transcript error. The x-axis represents the median difference in absolute log-fold error (Δ error = MAJEC error – competitor error). Negative values indicate lower quantification error for MAJEC. *Left:* Median error differences pooled across all timepoints, stratified by penalty type. *Right:* Median error differences for all expressed transcripts, stratified by biological timepoint and pooled overall. Horizontal lines indicate 95% bootstrap confidence intervals for the median difference. Percentages denote MAJEC’s superiority rate (the percentage of transcripts where MAJEC achieved a lower absolute error than the competitor, with ties distributed equally). Statistical significance was assessed using one-sided paired Wilcoxon signed-rank tests on non-zero differences (*** p < 0.001). **(C)** MAJEC transcript-level superiority rate against Salmon (blue) and RSEM (red), stratified by long-read ground truth expression bins (Counts Per Million, CPM). The dashed grey line at 0.5 represents performance parity; bars above this threshold indicate expression brackets where MAJEC outperforms the competitor for the majority of transcripts. Total transcript counts (*n*) per bin are annotated above the bars.

**Figure. S7.**
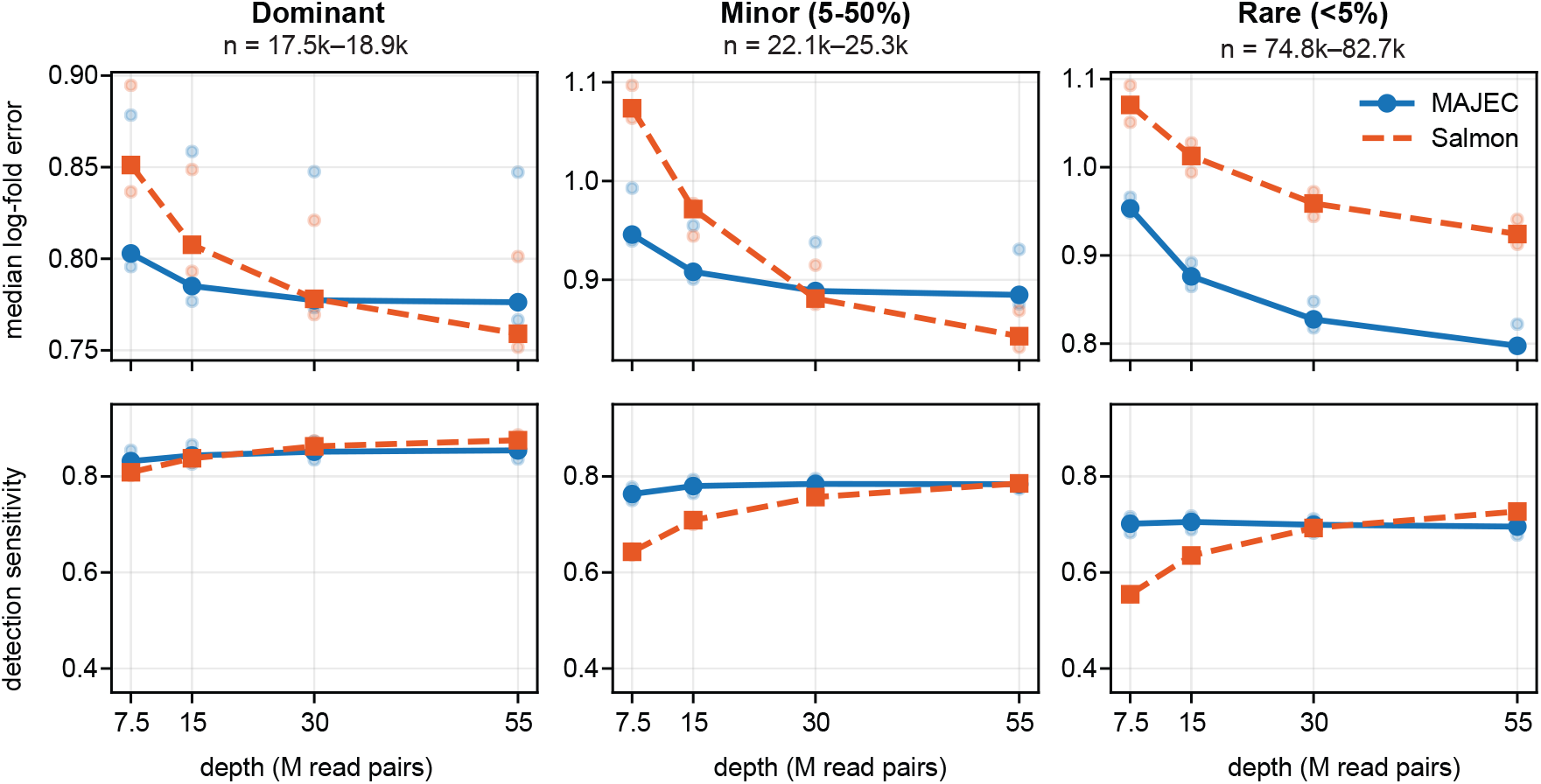
MAJEC versus Salmon accuracy and detection across sequencing depth in LongBench. Short reads from three cell lines (SHP-77, NCI-H2228, NCI-H146) were downsampled to 7.5, 15, 30, and 55 M read pairs and re-quantified with each tool, holding the matched long-read consensus ground truth fixed. Isoforms were classified from the long-read data into three disjoint per-gene classes: dominant (the highest-abundance isoform), minor (non-dominant isoforms contributing ≥5% of the gene’s long-read signal), and rare (non-dominant, <5%); n gives the per-line range of long-read-confirmed isoforms per class. Top row, median log-fold error; bottom row, detection sensitivity at a 0.1 CPM threshold (matching Fig 1E). Solid lines are medians across the three lines and faint points are individual lines. Lower error and higher sensitivity indicate better performance.

**Figure. S8.**
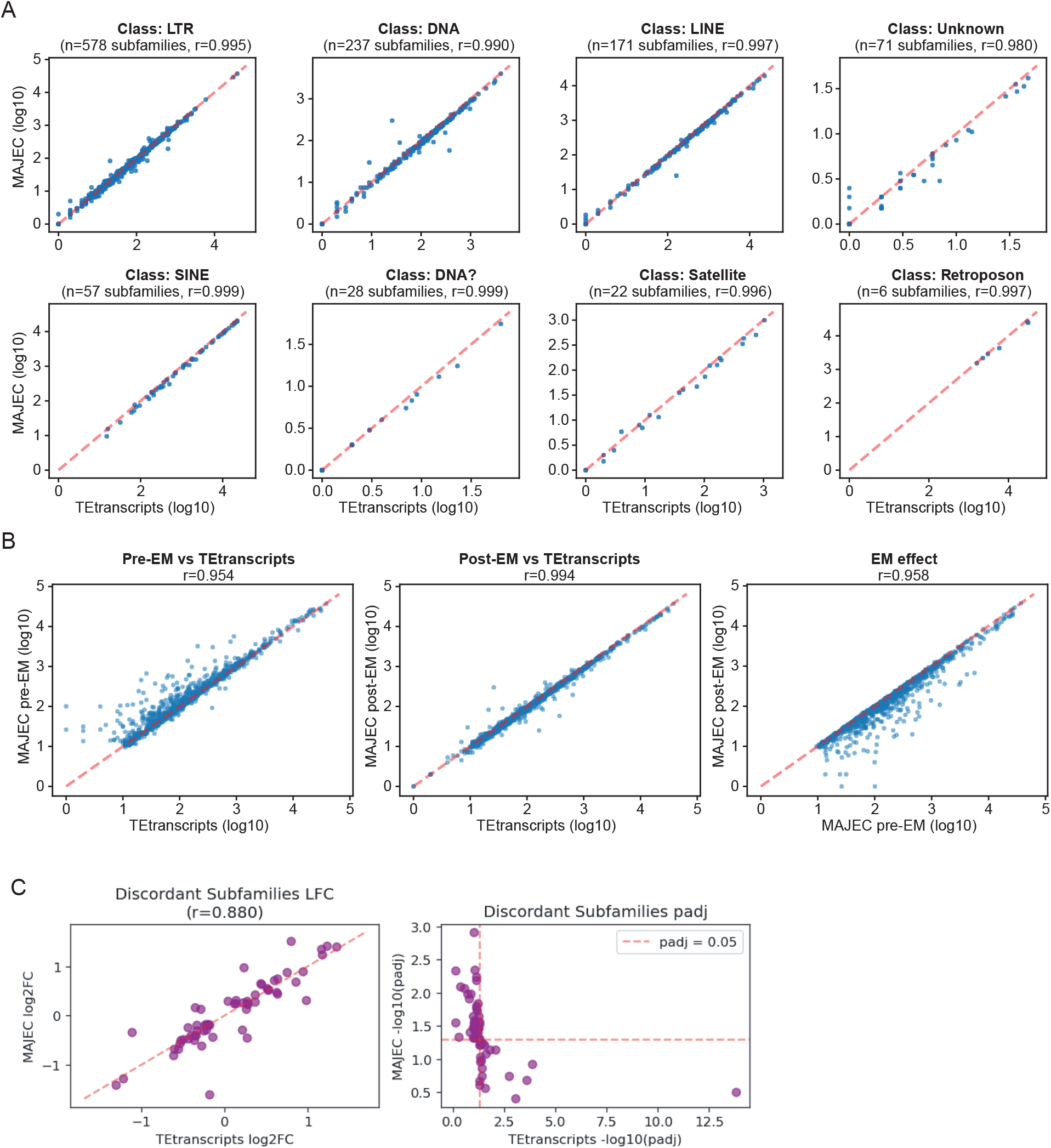
Comparison of MAJEC and TEtranscripts quantification and differential expression at the transposable element subfamily level. **(A)** Correlation of raw read counts between MAJEC and TEtranscripts across the top eight most abundant transposable element (TE) classes. Subfamily-level counts from a representative sample were log10(x+1) transformed. The number of subfamilies (n) and Pearson correlation coefficients (r) are indicated for each class. Dashed red lines represent the line of identity. **(B)** Impact of the Expectation-Maximization (EM) algorithm on MAJEC’s subfamily-level quantification. Scatter plots display log10(x+1) transformed counts comparing pre-EM MAJEC (calculated as initial unique alignments plus fractional multimappers) versus TEtranscripts (left), final post-EM MAJEC versus TEtranscripts (middle), and the direct effect of the EM algorithm on MAJEC counts (right). Application of the EM framework maximizes concordance with TEtranscripts (post-EM r = 0.994). **(C)** Left: Correlation of DEseq2 log2 fold change values for the 61 subfamilies called discordantly significant between the MAJEC and TEtranscripts datasets. Right: Scatter plot of significance for the 61 discordant subfamilies showing -log10 of the DEseq2 adjusted p-values. The discordant calls are still highly correlated and are generally close to the significance thresholds.

**Figure. S9.**
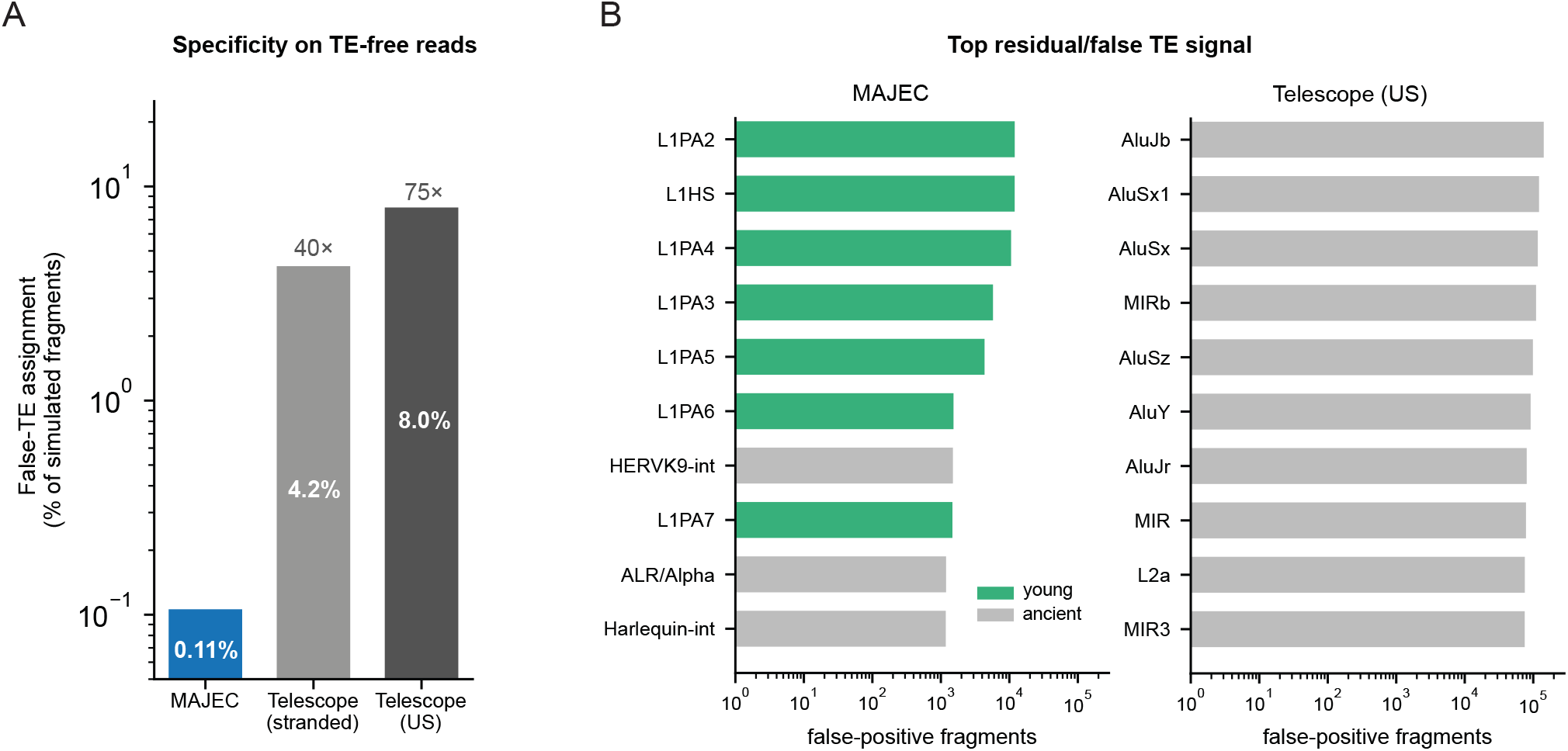
Specificity on a TE-free (gene-only) simulation. Paired-end reads (40M fragments, 150 bp) were simulated exclusively from GENCODE v44 protein-coding and lincRNA transcripts, with no TE sequence in the ground truth, so every read assigned to a TE locus is a false positive by construction. Telescope (US) is the released unstranded build; Telescope (stranded) is a development build. **(A)** False-TE assignment rate as a percentage of simulated fragments (log scale); fold values above the bars are relative to MAJEC. **(B)** TE subfamilies contributing the most false-positive fragments for MAJEC (left) and released Telescope (right) on a shared log scale, colored by family age (young L1, comprising L1HS and L1PA; ancient, all other families). Values are from one simulated library (sample_01).

**Figure. S10.**
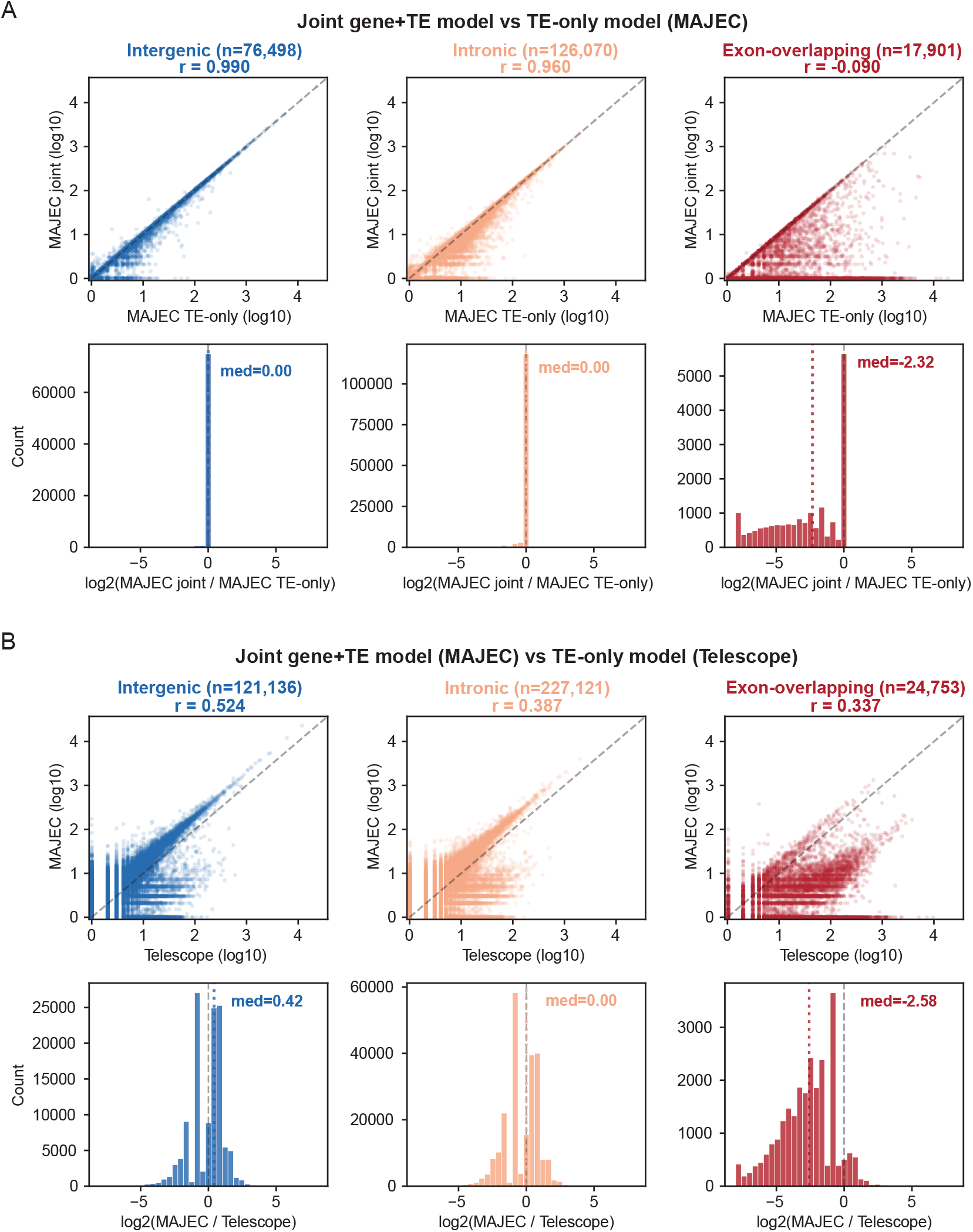
Influence of gene overlap on locus-level transposable element quantification in joint versus TE-only models. Transposable element (TE) loci were classified based on their genomic context relative to protein-coding genes: intergenic (blue), intronic (orange), or exon-overlapping (red). **(A)** Comparison of MAJEC run in its standard joint gene+TE mode versus a restricted TE-only mode (where gene annotations were excluded). *Top row:* Scatter plots of log10(x+1) transformed locus counts. The dashed grey line represents the line of identity. Pearson correlation coefficients (r) and the total number of loci with a count >= 1 in either model (n) are indicated. *Bottom row:* Histograms displaying the distribution of log2 fold changes (MAJEC joint / MAJEC TE-only) for expressed loci. The dashed grey line indicates a log2 fold change of zero (no difference), while the heavy dotted colored line marks the median difference (med). **(B)** Comparison of MAJEC (joint gene+TE model) against Telescope (an independent TE-only locus quantification tool). *Top row:* Scatter plots of log10(x+1) transformed locus counts across the three overlap classes. *Bottom row:* Histograms displaying the distribution of log2 fold changes (MAJEC / Telescope). As with the internal MAJEC comparison, exon-overlapping TEs show severe systemic disagreement (median log2FC = −2.58), highlighting the massive over-counting of genic reads by TE-only models when the joint feature space is ignored.

**Fig. S11.**
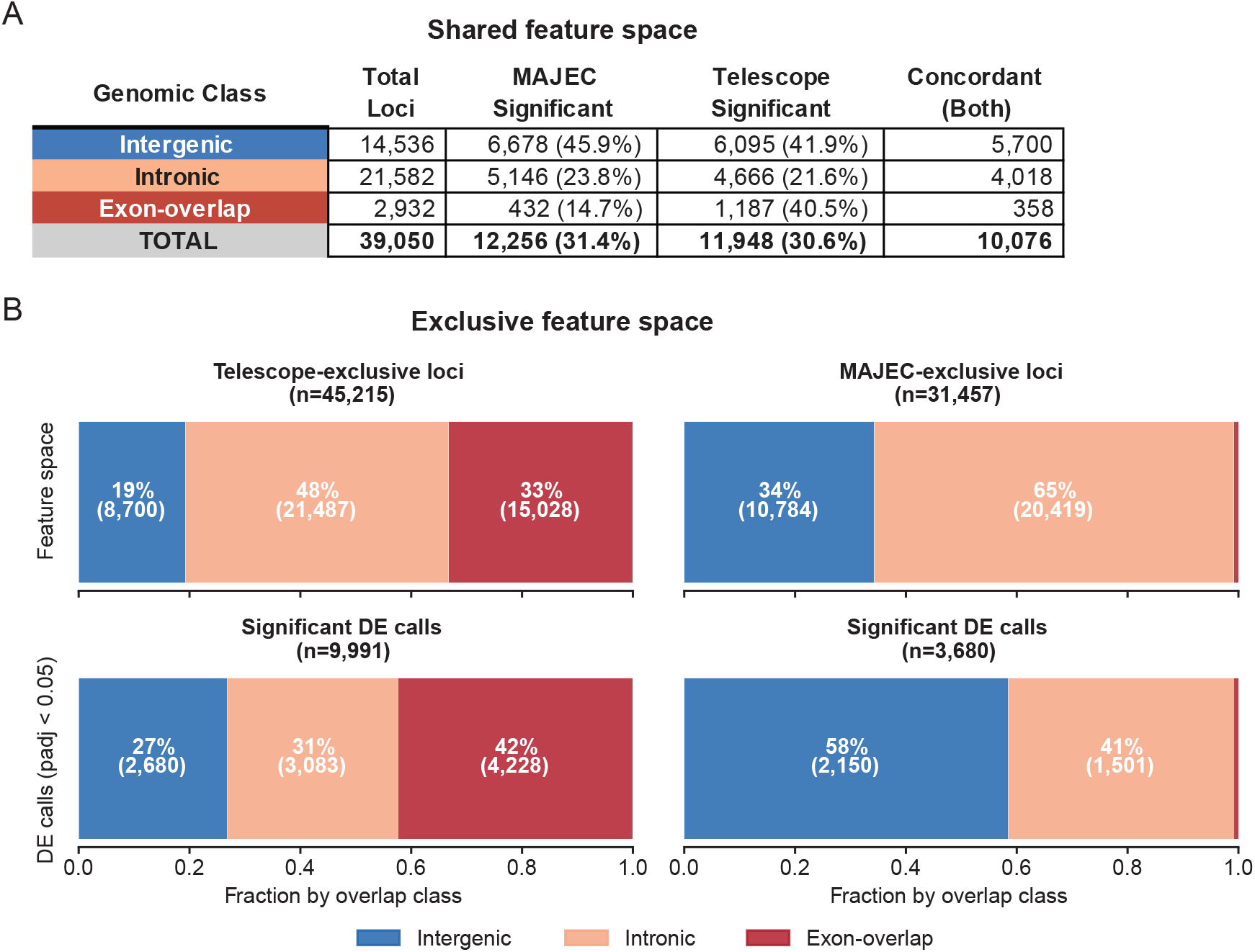
Genomic context of shared and tool-exclusive transposable element differential expression. **(A)** Summary of differential expression (DE) calls within the shared feature space (39,050 TE loci that passed the minimum count threshold in both MAJEC and Telescope). The table displays the total number of loci per genomic class (intergenic, intronic, and exon-overlapping) alongside the number and percentage of significant DE calls (adjusted p-value < 0.05) made by each tool independently, as well as the concordant calls identified by both. **(B)** Composition of the exclusive feature space, defined as TE loci that passed the minimum expression filter in one tool but not the other. *Top row:* The proportion of all tool-exclusive loci assigned to each genomic overlap class for Telescope (left) and MAJEC (right). *Bottom row:* The proportion of significant DE calls (padj < 0.05) within those exclusive feature spaces. Total locus counts (*n*) and percentages are annotated within the bars. Consistent with the lack of a joint feature space, Telescope’s DE calls are heavily enriched for exon-overlapping and intronic elements, characteristic of host gene misattribution. In contrast, MAJEC suppresses these exonic counts, yielding an exclusive DE profile highly enriched for intergenic loci (58%) that represent unambiguous, autonomous TE transcription.

**Fig. S12.**
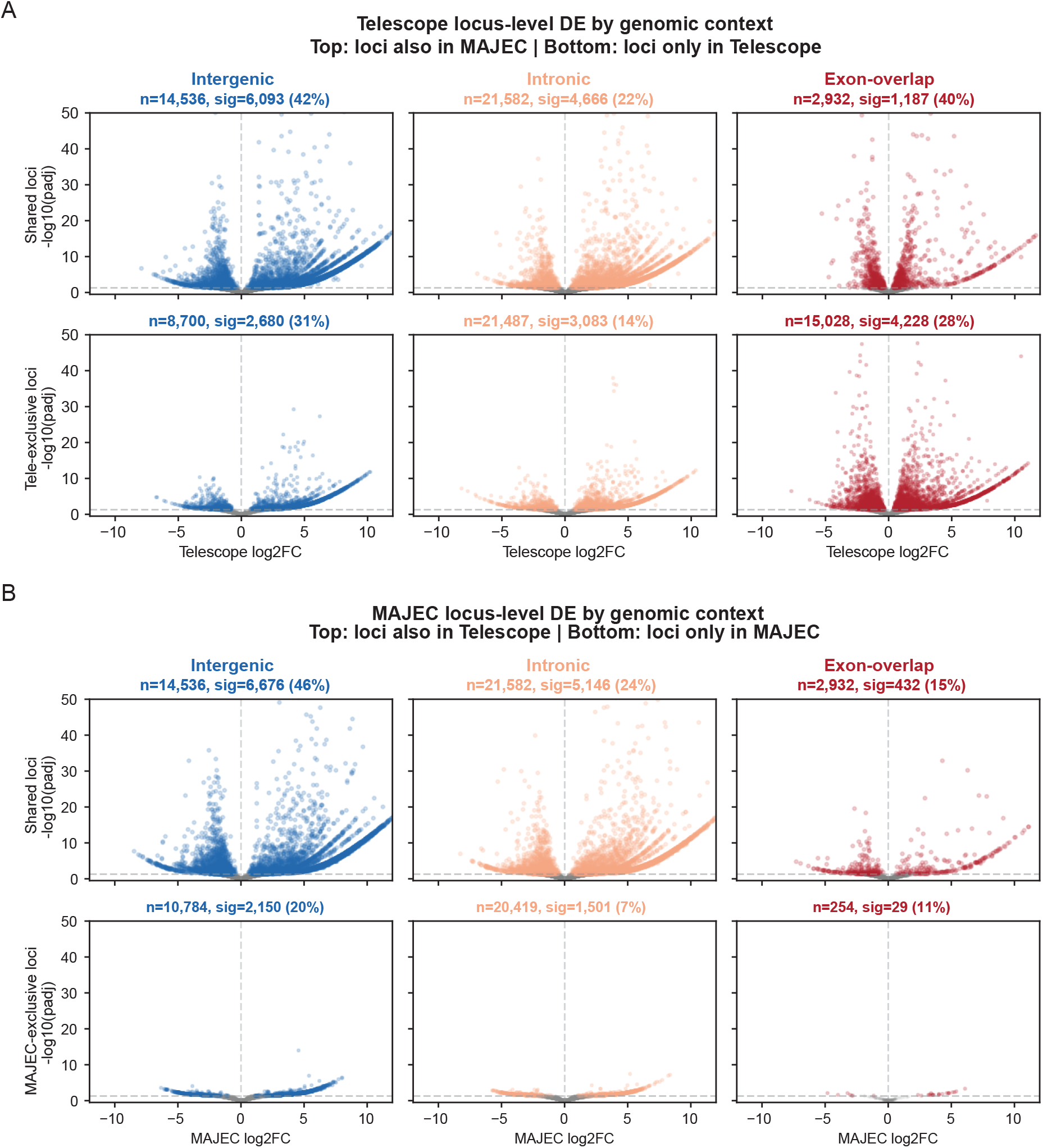
Volcano plots of locus-level differential expression stratified by genomic context and shared versus exclusive feature spaces. **(A)** Differential expression of transposable element (TE) loci quantified by Telescope. Plots display the log2 fold change (x-axis) against the -log10 adjusted p-value (y-axis). Columns separate loci based on their genomic context relative to protein-coding genes: intergenic (blue), intronic (orange), and exon-overlapping (red). The top row displays loci within the shared feature space (passing minimum count thresholds in both Telescope and MAJEC), while the bottom row displays Telescope-exclusive loci. The horizontal dashed grey line indicates the significance threshold (padj = 0.05). Total locus counts (n) and the number/percentage of significant DE calls (sig) are annotated above each plot. **(B)** Equivalent differential expression volcano plots for TE loci quantified by MAJEC, split into shared loci (top row) and MAJEC-exclusive loci (bottom row). Notably, Telescope generates a massive volume of highly significant, large-magnitude fold changes within its exclusive intronic and exon-overlapping feature spaces (A, bottom row). In contrast, the MAJEC-exclusive feature space exhibits a starkly flattened significance profile (B, bottom row), visually demonstrating that MAJEC’s joint EM framework successfully suppresses the spurious, high-significance fold changes driven by genic read misattribution.

**Fig. S13.**
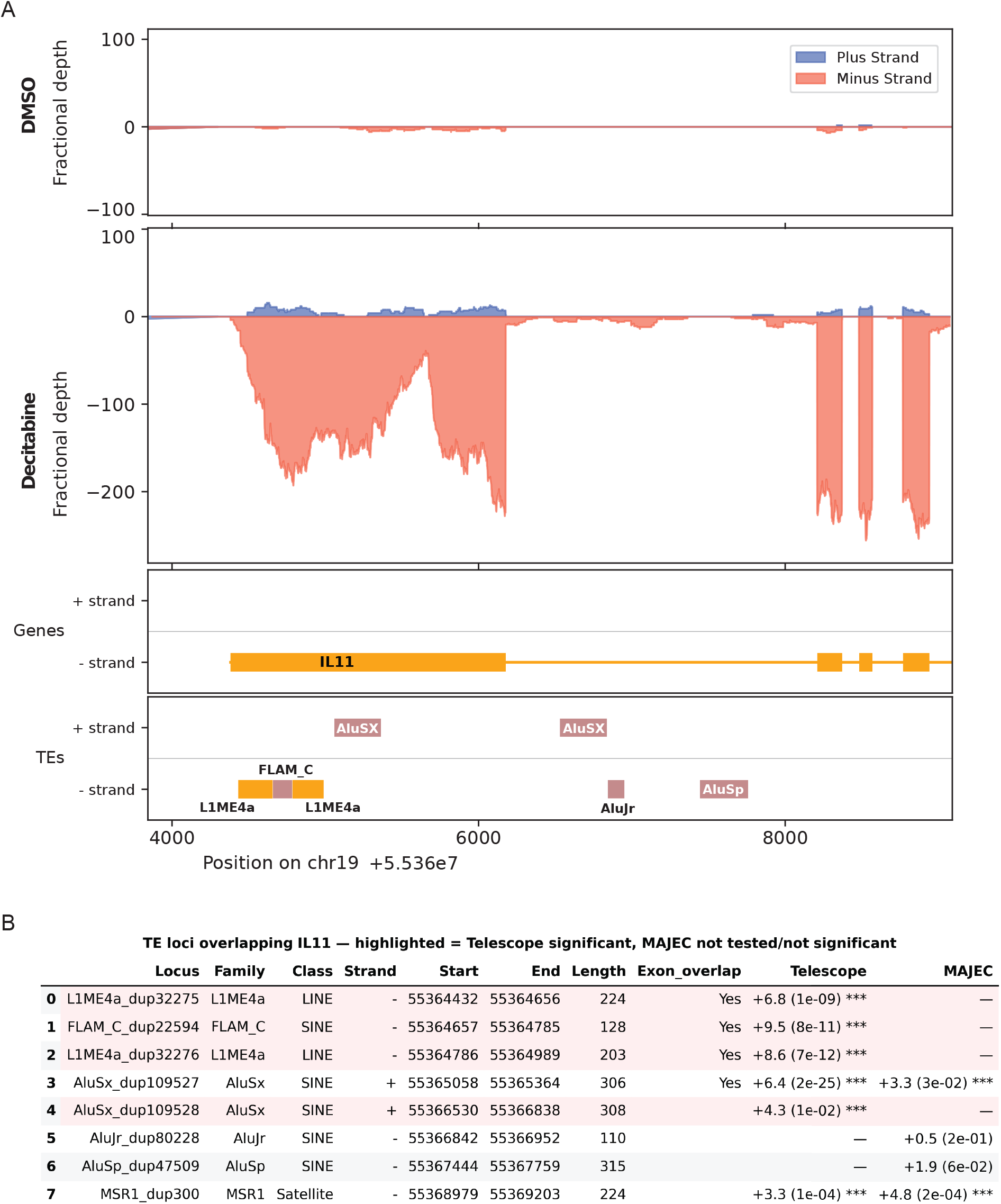
Validation of Telescope false reactivation artifacts at the *IL11* locus following decitabine treatment. **(A)** Stranded fractional read coverage across the *IL11* gene locus (chr19) in DMSO control (top) and decitabine treatment (middle) conditions. Plus-strand (blue) and minus-strand (red) coverage are plotted above gene and transposable element (TE) annotation tracks (bottom). Decitabine treatment induces massive minus-strand transcription that precisely matches the *IL11* exon-intron structure, with no independent transcriptional peaks isolated over the numerous embedded TE annotations. **(B)** Differential expression summary for all TE loci overlapping the *IL11* gene body. Rows highlighted in red represent loci that Telescope incorrectly reported as significantly upregulated (padj < 0.05) but which MAJEC correctly recognized as genic signal (either not tested due to insufficient counts or not significant). The log2 fold change and adjusted p-values are provided for each tool. Telescope systemically misattributes the drug-induced upregulation of the *IL11* gene to the embedded TE fragments.

**Table S1.** Misattribution of TE reactivation signals to host genes by hierarchical quantification. Top candidate loci demonstrating signal misattribution when using TEtranscripts count matrices. Candidates were selected based on identifying strong TE-driven upregulation (TE LFC > 1) that exceeded the host gene’s signal magnitude. **MAJEC TE LFC** and **MAJEC Gene LFC** (along with corresponding p-adj values) reflect differential expression metrics calculated by DESeq2 using the probabilistically resolved count estimates from MAJEC’s joint EM model. **TET Gene LFC** and **TET p-adj** reflect the corresponding DESeq2 results when using the count matrix generated by TEtranscripts. **LFC Inflation** is calculated as the difference between the TET Gene LFC and the MAJEC Gene LFC, representing the magnitude of false-positive gene signal generated by absorbed TE reads. Top misattributed loci are highlighted in red (Inflation > 1.0).

| TE Locus | Subfamily | Host Gene | MAJEC<br>TE LFC | MAJEC<br>Gene LFC | MAJEC p-adj | TET Gene<br>LFC | TET p-adj | LFC<br>Inflation |
| --- | --- | --- | --- | --- | --- | --- | --- | --- |
| LTR1A2_dup786 | LTR1A2 | LINC02575 | 4.29 | 0.75 | 2.96E-03 | 8.43 | 1.51E-09 | 7.69 |
| LTR1A2_dup787 | LTR1A2 | LINC02575 | 4.28 | 0.75 | 2.96E-03 | 8.43 | 1.51E-09 | 7.69 |
| LTR12C_dup2304 | LTR12C | LINC01467 | 4.76 | 0.23 | 2.07E-01 | 6.86 | 2.39E-06 | 6.63 |
| L1PA7_dup4347 | L1PA7 | LINC01949 | 6.04 | 0.24 | 2.00E-01 | 5.35 | 2.64E-09 | 5.11 |
| HERVH-int_dup353 | HERVH-int | LINC02474 | 5.81 | 0.37 | 2.65E-01 | 5.15 | 6.82E-04 | 4.78 |
| LTR12C_dup1283 | LTR12C | LINC01447 | 3.62 | 0.21 | 5.13E-01 | 4.7 | 2.14E-03 | 4.49 |
| L1PB1_dup3753 | L1PB1 | LINC02492 | 5.65 | 0.33 | 3.61E-01 | 4.48 | 2.93E-03 | 4.15 |
| LTR12_dup159 | LTR12_ | CTD-3080P12.3 | 3.05 | 0.47 | 2.15E-01 | 1.85 | 4.31E-02 | 1.38 |
| HAL1_dup16670 | HAL1 | LINC00592 | 2.57 | 0.53 | 3.98E-01 | 1.78 | 6.65E-06 | 1.25 |
| HERVIP10F-int_dup195 | HERVIP10F-int | LINC00431 | 4.41 | 2.65 | 1.39E-17 | 3.48 | 2.37E-47 | 0.83 |
| L1PA15_dup1663 | L1PA15 | LOC101928882 | 4.45 | 2.45 | 1.72E-02 | 3.15 | 4.71E-03 | 0.7 |
| L1MC4_dup7359 | L1MC4 | LINC02228 | 6.25 | 2.76 | 4.71E-15 | 3.29 | 4.15E-19 | 0.52 |
| HERVH-int_dup5748 | HERVH-int | TPTEP1 | 5.28 | 3.52 | 1.24E-09 | 4.03 | 8.31E-09 | 0.51 |
| L1PA3_dup10297 | L1PA3 | ZNF66 | 6.76 | 5.38 | 4.38E-11 | 5.64 | 1.07E-07 | 0.25 |
| SVA_D_dup880 | SVA_D | MRGPRX3 | 4.56 | 0.99 | 7.20E-02 | 1.23 | 3.71E-02 | 0.24 |
| HERVIP10B3-int_dup180 | HERVIP10B3-int | LOC101928595 | 4.4 | 2.52 | 1.80E-27 | 2.57 | 6.41E-23 | 0.05 |
| L2_dup896 | L2 | CDC20 | 2.94 | 1.93 | 9.91E-157 | 1.92 | 9.40E-151 | 0 |
| MIRc_dup66682 | MIRc | FOSL1 | 4.25 | 3.03 | 0.00E+00 | 3.03 | 0.00E+00 | 0 |

**Table S2.**
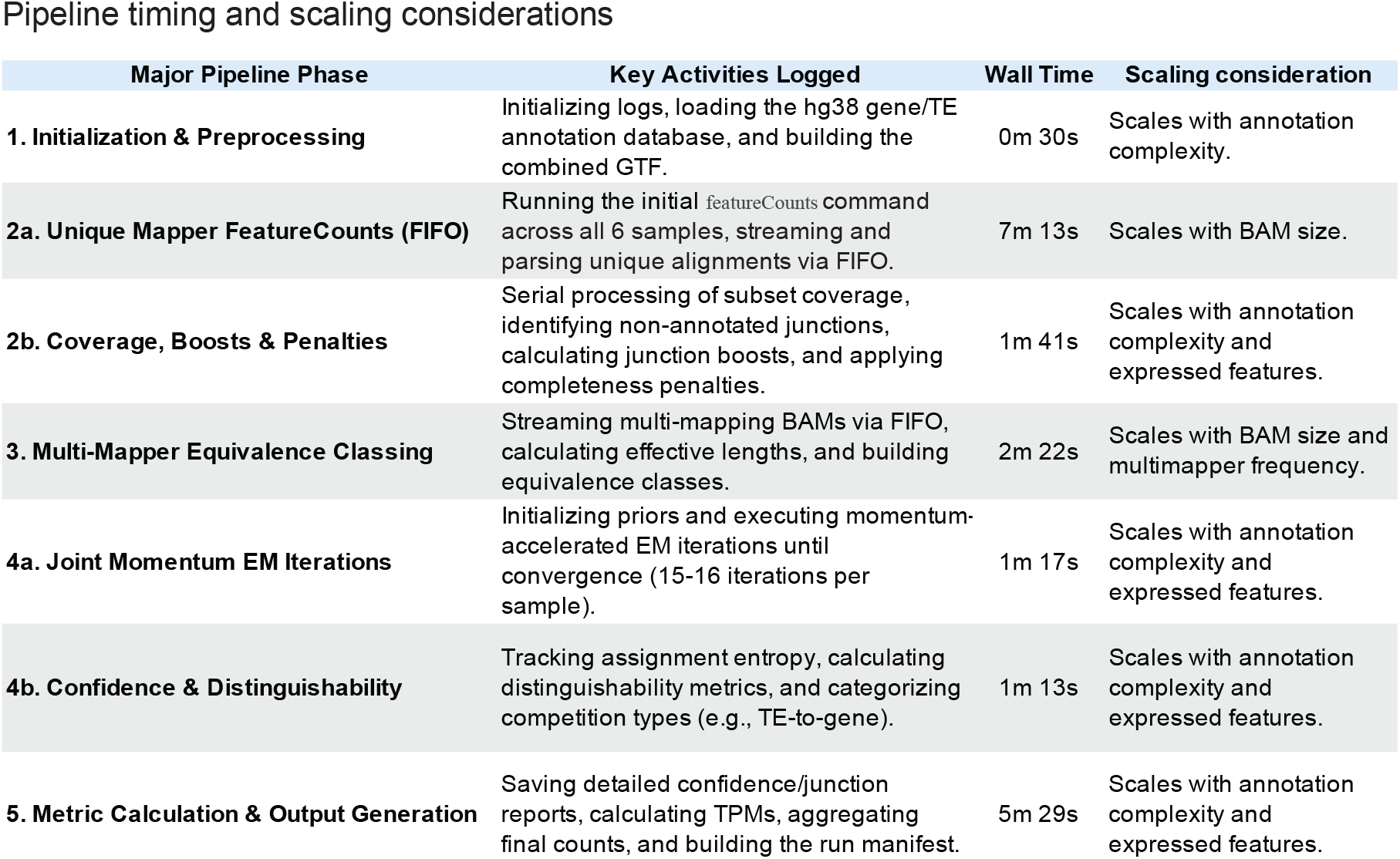
Detailed timing and computational scaling profile of the MAJEC pipeline. Breakdown of wall time and scaling factors for each major phase of the MAJEC algorithm. Times reported reflect the joint processing of six RNA-seq samples (averaging **∼53.5 million reads per sample**) on an Intel Xeon HPC cluster, totaling approximately 20 minutes. The underlying feature space is highly complex, comprising the full human reference transcriptome combined with RepeatMasker TE annotations for a total of **4,770,531 annotated features**. The execution time is dominated by BAM file I/O and alignment parsing via featureCounts (Phases 2a and 3), which scale linearly with input read depth. In contrast, the core mathematical modeling—including the joint momentum-accelerated EM algorithm (Phase 4a) and distinguishability metric calculations (Phase 4b)—is highly optimized. Despite tracking between 250,000 and 350,000 actively expressed features per sample, these joint modeling steps complete in under three minutes combined.

